# Benchmarking AlphaFold and related deep learning approaches for modeling antibody and TCR antigen recognition

**DOI:** 10.64898/2026.07.04.736425

**Authors:** Rui Yin, Shayana Saravanakumar, Shu Yuan Shi, Minjae Park, Valerie Lin, Jessica Lee, Melyssa Cheung, Nathaniel Felbinger, Sivan Kaufman, Maya Eisenberg, Brian G. Pierce

## Abstract

Determining the structural basis of antigen recognition by antibodies and T cell receptors (TCRs) provides critical insights into effective immune targeting and can inform design of biotherapeutics and vaccines. Accurate computational modeling of antibodies and TCRs in complex with their targets poses a major challenge for predictive methods, including AlphaFold, which is generally accurate for modeling protein complexes but has shown limited success for immune recognition. In this study we assessed the performance of AlphaFold2, AlphaFold3, increased sampling protocols, and related deep learning methods for modeling antibody-protein, antibody-peptide, and TCR-peptide-major histocompatibility complex (pMHC) recognition. We show that increased sampling and AlphaFold3 generally improve performance relative to default sampling and AlphaFold2, however predictive accuracy and improvement levels varied considerably among interface classes, with antibody-peptide complexes representing a challenge despite their small antigen size. Comparing per-case success across methods showed some complementarity, indicating opportunities for increased success through model pooling approaches, for instance increasing antibody-peptide near-native success from 41% to 59%. Analysis of AlphaFold confidence scores and modeling of a noncanonical complex provided further insights into predictive performance. These results highlight considerations for predictive antibody and TCR complex modeling efforts, while revealing key distinctions among protocols, scoring, and immune complex classes.

## Introduction

Antibodies and T cell receptors (TCRs) are key components of immunity, responsible for specific recognition of proteins, peptides, and other molecular antigens from an immense array of viruses and pathogens, thereby initiating B cell and T cell immune responses^1^. They also represent important and growing classes of biotherapeutics^2–4^. High resolution structures of antibodies and TCRs in complex with their targets provide critical insights into effective and aberrant immune responses, for instance revealing targeting strategies of broadly neutralizing antibodies^5–8^, autoimmune TCR recognition of self-antigens^9, 10^, recognition of cancer neoantigens by TCRs^11^, and the mechanistic basis for viral escape^12–14^. Additionally, such structures provide the means to perform structure-based design of antibodies and TCRs to improve their targeting and other properties^15–18^, as well as design of vaccine antigens to elicit more optimal immune responses^19–21^. Despite their importance, the number of experimentally determined antibody and TCR complex structures remains limited due to the cost, time, and technical challenges of experimental structure determination, particularly in light of the scale of immune repertoires comprising millions of distinct antibodies and TCRs.

To address this need, there has been growing interest in the development and use of computational methods to accurately model structures of antibodies and TCRs in complex with their targets. Previously developed complex prediction methods using docking approaches showed limited success in generating near-native complex models^22, 23^, in part due to challenges with modeling conformational flexibility in the complementarity determining region (CDR) loops that engage the antigen, as well as flexibility in the antigens themselves, during the docking process. Additionally, while docking approaches can output near-native models of antibody-antigen complexes in large sets of models for a complex, they are often not ranked highly, leading to incorrect top-ranked models in many cases^22^. Due to TCRs binding their peptide and major histocompatibility complex (pMHC) targets with a generally conserved diagonal mode, some have employed homology modeling based approaches to model TCR-pMHC complexes^24, 25^, yet the diversity of TCR binding modes and CDR loop conformations^23, 26, 27^ limits the accuracy of such approaches for modeling of unseen TCRs and their interactions^28^.

The development of AlphaFold2^29^, AlphaFold3^30^, and related deep learning structure prediction methods has provided an avenue to accurately model antibody-antigen and TCR-pMHC complexes from sequence. Such deep learning-based “fold-and-dock” complex assembly was found to outperform previously developed modeling approaches for antibody-antigen^31^ and TCR-pMHC ^28^ complexes. However, high accuracy near-native models were still not generated for the majority of antibody and TCR complexes with those approaches^28, 31^. This limitation is likely due to the lack of useful co-evolutionary signal across antibody-antigen and TCR-pMHC interfaces in the AlphaFold input multiple sequence alignments (MSAs)^31, 32^, as well as insufficient numbers of distinct antibody-antigen and TCR-pMHC complex structures in the Protein Data Bank (PDB) for AlphaFold training, on the order of several thousand antibody-antigen^33^ and several hundred TCR-pMHC^34^ complex structures. To address limitations in AlphaFold complex modeling accuracy, several groups have found that massive sampling in AlphaFold2, entailing pooling large sets of AlphaFold2-based models for a complex and ranking by AlphaFold2 confidence score, led to improved accuracy for antibody-antigen modeling in recent Critical Assessment of Structural Prediction (CASP) and Critical Assessment of Predicted Interactions (CAPRI) blind structure prediction rounds^35–37^ and in benchmarking^31^. AlphaFold3, which uses a diffusion model along with a more general all-atom representation of molecules, was found by its developers to outperform AlphaFold2 for antibody-antigen modeling accuracy^30^, and this was corroborated by initial benchmarking by others^38^. Notably, its reported success on antibody-antigen interfaces was dependent on the number of models generated per complex, with noted success increasing with up to 5000 models per complex^30^. As with AlphaFold2 massive sampling, this increased AlphaFold3 sampling requires substantial computational resources, while the observed success for high accuracy models at that level of sampling was higher than for AlphaFold2 but still fewer than half of tested complexes (∼30%)^30^. AlphaFold3-like all-atom deep learning modeling methods Chai-1^39^ and Boltz-1^40^ are available open source and for commercial use, unlike AlphaFold3, but initial benchmarking has shown that those methods have lower modeling accuracy for antibody-antigen complexes versus AlphaFold3^38^.

Several limitations in previous studies have resulted in open questions regarding the utility, practical considerations, and optimal use of deep learning approaches for modeling adaptive immune recognition. Notably, the accuracy of AlphaFold modeling for antibody-peptide complexes, which represent an important and therapeutically relevant class of complexes with distinct interface structural features from antibody-protein complexes^41^, has not been systematically evaluated outside of anecdotal benchmarking of limited nanobody-peptide^42^ or antibody-peptide^43^ complexes, and performance on individual CAPRI targets^44^. Due to the intrinsic flexibility of peptides and antibody CDR loops^41^, antibody–peptide complexes present a challenge for docking-based approaches, and they represent a compelling target for structure prediction from sequence using AlphaFold. In addition, TCR-pMHC structure prediction has not been assessed in detail for AlphaFold3 and increased sampling approaches, other than recent AlphaFold3 benchmarking in the absence of increased sampling^45^ and anecdotal modeling of newly determined complex structures^46, 47^.

In this study we present a comprehensive analysis of AlphaFold2, AlphaFold3, and AlphaFold3-related deep learning methods for antibody-protein, antibody-peptide, and TCR-pMHC complex structure prediction. We assessed the impact of increased sampling approaches on modeling success for each class of interface, as well as the association of confidence scores with modeling accuracy levels. Additionally, we identified highly variable modeling performance across approaches and interface classes, as well as the utility of confidence scores for identification of highly accurate models. While AlphaFold3 showed highest performance of the assessed methods, there was complementarity in modeling accuracy of complexes from other approaches, indicating that pooled approaches can provide the means for improving upon its still limited success. This highlights important steps toward accurate generalizable prediction of unseen antibody-antigen and TCR-pMHC complex structures from sequence, applicable for modeling of structurally uncharacterized interfaces as well as antibody and TCR design pipelines, indicating directions for future developments.

## Results

### AlphaFold3 and increased sampling accuracy for immune complex classes

To assess and compare AlphaFold2 (AF2) and AlphaFold3 (AF3) accuracy for antibody-protein, antibody-peptide, and TCR-pMHC complexes, we assembled sets of nonredundant complex structures from the Protein Data Bank (PDB) released after the AF2 and AF3 training date cutoff (9/30/2021) (**Table S1**, **Table S2**, **Table S3**). This led to 80 antibody-antigen complexes, 41 antibody-peptide complexes, and 20 TCR-pMHC complexes for benchmarking, with the set of TCR-pMHC cases corresponding to those used in a previous assessment^28^.

We first tested antibody-protein modeling performance in AF2 and AF3 with default and increased sampling. Four protocols were run on the benchmark set: 1) AF2 (v2.3 multimer model)^48^ with default parameters (25 structures per complex), 2) AF2 massive sampling, which generated a total of 8000 structures per complex using varying parameters and trained AF2 models (v2.1, v2.2, v2.3), as detailed in the Methods, 3) AF3 with default parameters (5 structures per complex), and 4) AF3 with increased sampling, corresponding to 200 random seeds and 1000 structures per complex. For each method, structural models for a complex were ranked by confidence score, and top-ranked models were assessed for accuracy based on CAPRI criteria^49^ to obtain success rates on the benchmark set (**Fig. 1**). Top-ranked model success rate (T1) for default AF2 was approximately 30% for near-native complexes (medium or high CAPRI accuracy), which increased to nearly 50% for the AF2 massive sampling approach. These accuracy levels are similar to our previous benchmarking on a smaller set of antibody-protein complexes (N=37) and a moderately different massive sampling protocol^31^. AF3 performance was found to be higher than AF2 performance when comparing the respective default and increased sampling protocols, with T1 near-native (medium/high) accuracy success of approximately 38% and 60% for AF3 default and AF3 increased sampling, respectively. This increase in performance for antibody-protein antigen modeling for AF3 versus AF2 corroborates the improvement noted by the AlphaFold developers ^30^ and benchmarking by others^38^.

**Figure 1.**
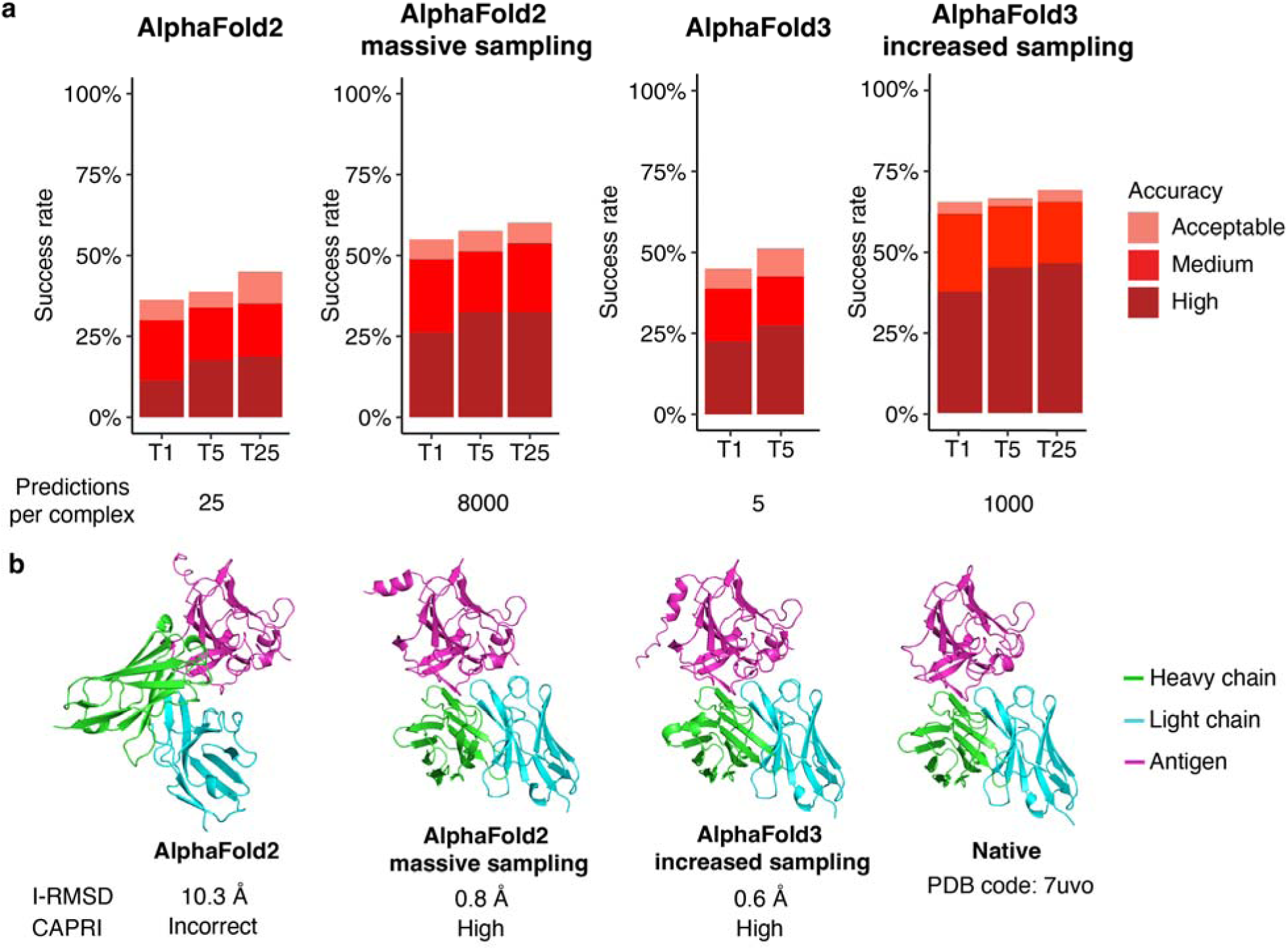
AlphaFold success for antibody-protein complex modeling. (a) Success rates for AlphaFold2 and AlphaFold3 with default and massive/increased sampling protocols for top-ranked (T1), top 5 (T5) and top 25 (T25, when available) ranked models per complex (80 test cases). Number of total models generated per complex are shown at bottom. Bars are colored by CAPRI accuracy levels, as shown on right. (b) Example top-ranked models of antibody-protein complex, with native complex structure (PDB code 7uvo) on right. Antibody and antigen chains are colored as indicated on right. Interface RMSD (I-RMSD) from native complex and CAPRI accuracy level for each model are shown at bottom.

We also assessed AF2 and AF3 performance for the antibody-peptide complex benchmark set, using default and increased sampling methods for each protocol. A moderately reduced increased sampling protocol was used for AF2 versus antibody-protein complexes, generating 1500 structures per complex, while for AF3 increased sampling 1000 structures were generated. Assessed antibody-peptide modeling accuracy (**Fig. 2**) showed lower performance in AF2 and AF3 versus antibody-protein complexes. Top-ranked model high accuracy success was approximately 12% for default AF2 and AF3, with moderate improvements from increased sampling (20% high accuracy success for each). The near-native medium/high success was approximately equal for AF2 and AF3 increased sampling (35-40%). In comparison with antibody-protein complexes, overall AF2 and AF3 accuracies were lower, no substantial improvement was observed from AF3 with respect to AF2, and increased sampling led to more moderate gains in success for antibody-peptide complexes. This highlights notable differences between these complex classes and an increased challenge from antibody-peptide complexes for deep learning modeling, despite smaller antigen size.

**Figure 2.**
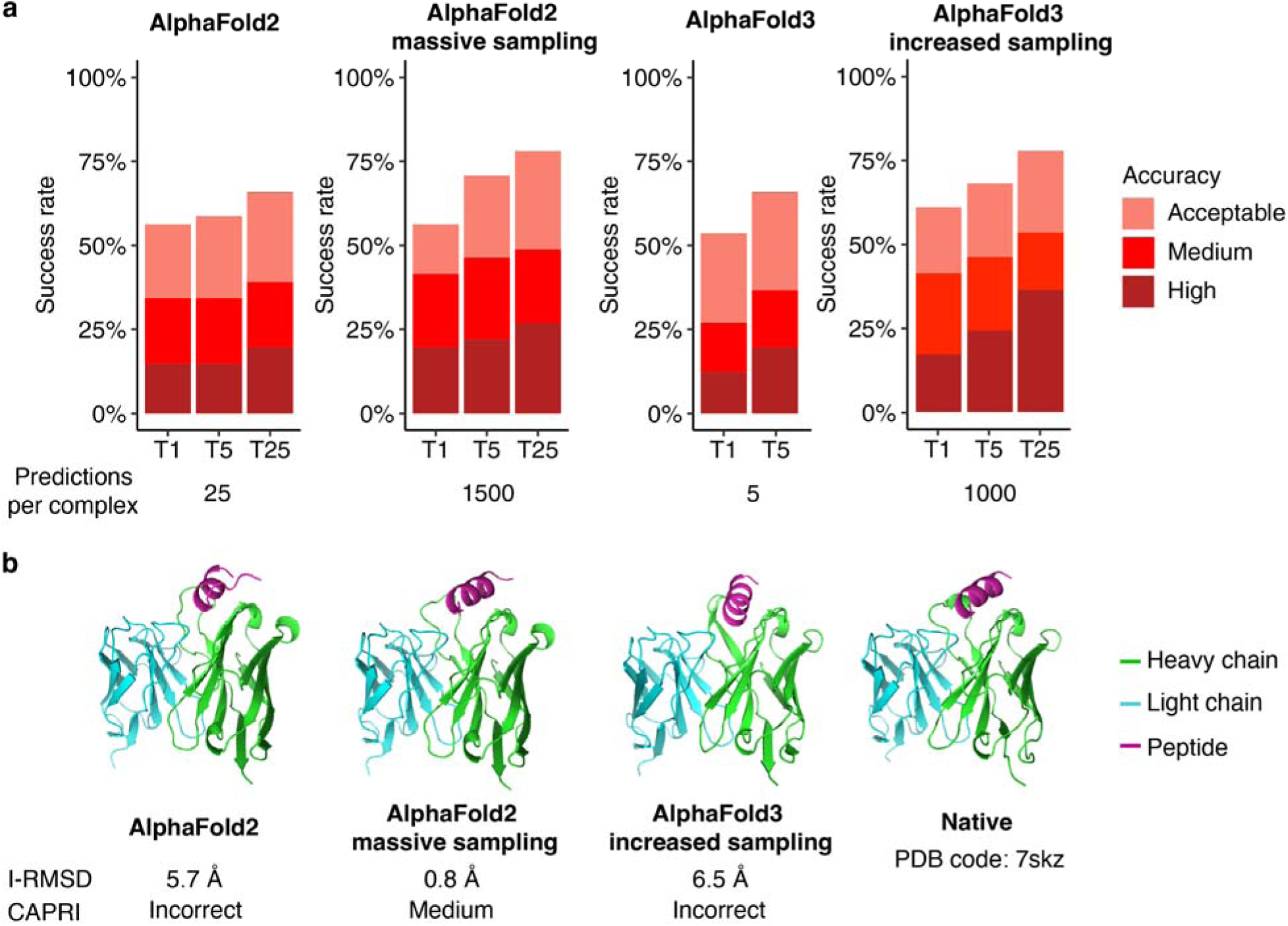
AlphaFold success for antibody-peptide complex modeling. (a) Success rates for AlphaFold2 and AlphaFold3 with default and massive/increased sampling protocols for top-ranked (T1), top 5 (T5) and top 25 (T25, when available) ranked models per complex (41 test cases). Number of total models generated per complex are shown at bottom. Bars are colored by CAPRI accuracy levels, as shown on right. (b) Example top-ranked models of antibody-peptide complex, with native complex structure (PDB code 7skz) on right. Antibody and peptide chains are colored as indicated on right. Interface RMSD (I-RMSD) from native complex and CAPRI accuracy level for each model are shown at bottom.

TCR-pMHC complex modeling performance was benchmarked for baseline and increased sampling AF2 and AF3 protocols (**Fig. 3**), using a set of 14 Class I and 6 Class II MHC complexes (**Table S3**). For AF2, in addition to the default protocol, we tested TCRmodel2 which is an optimized adaptation of AF2 that uses the same deep learning model and general pipeline, but has a TCR- and MHC-focused MSA database and optimized TCR chain and pMHC template structure identification^28^. In contrast with antibody-protein and antibody-peptide complexes, all protocols had TCR-pMHC top-ranked model (T1) success greater than 60% for medium or higher accuracy models (**Fig. 3**). However, generating high accuracy models across the benchmark set still posed a major challenge, and coupled with the need for high model accuracy to show detailed TCR-pMHC interface features and contacts^46^, high accuracy success was prioritized for TCR-pMHC cases. The default TCRmodel2 success rates modestly surpassed those of AF2 (as previously observed^28^), while performing massive sampling in TCRmodel2 (1500 structures generated per complex) led to a notable improvement of high accuracy success for top-ranked (T1) and top 5 ranked (T5) models versus default TCRmodel2 and AlphaFold2. Additionally, TCRmodel2 with massive sampling exhibited a 100% T5 success for medium or better accuracy models, and an over 50% T25 success rate for high accuracy models. The default AF3 TCR-pMHC top-ranked high accuracy success rate was higher than or comparable to default AF2 and TCRmodel2. Increased sampling in AF3 led to some improvement in performance versus default single-seed performance for both high and medium/high success, with T1 success rates matching those of TCRmodel2 with massive sampling. Overall, AF3 and increased sampling yielded some success improvements for TCR-pMHC modeling with respect to baseline AF2 and default single-seed protocols, although the improvements were generally less pronounced than for antibody-protein complexes (**Fig. 1**).

**Figure 3.**
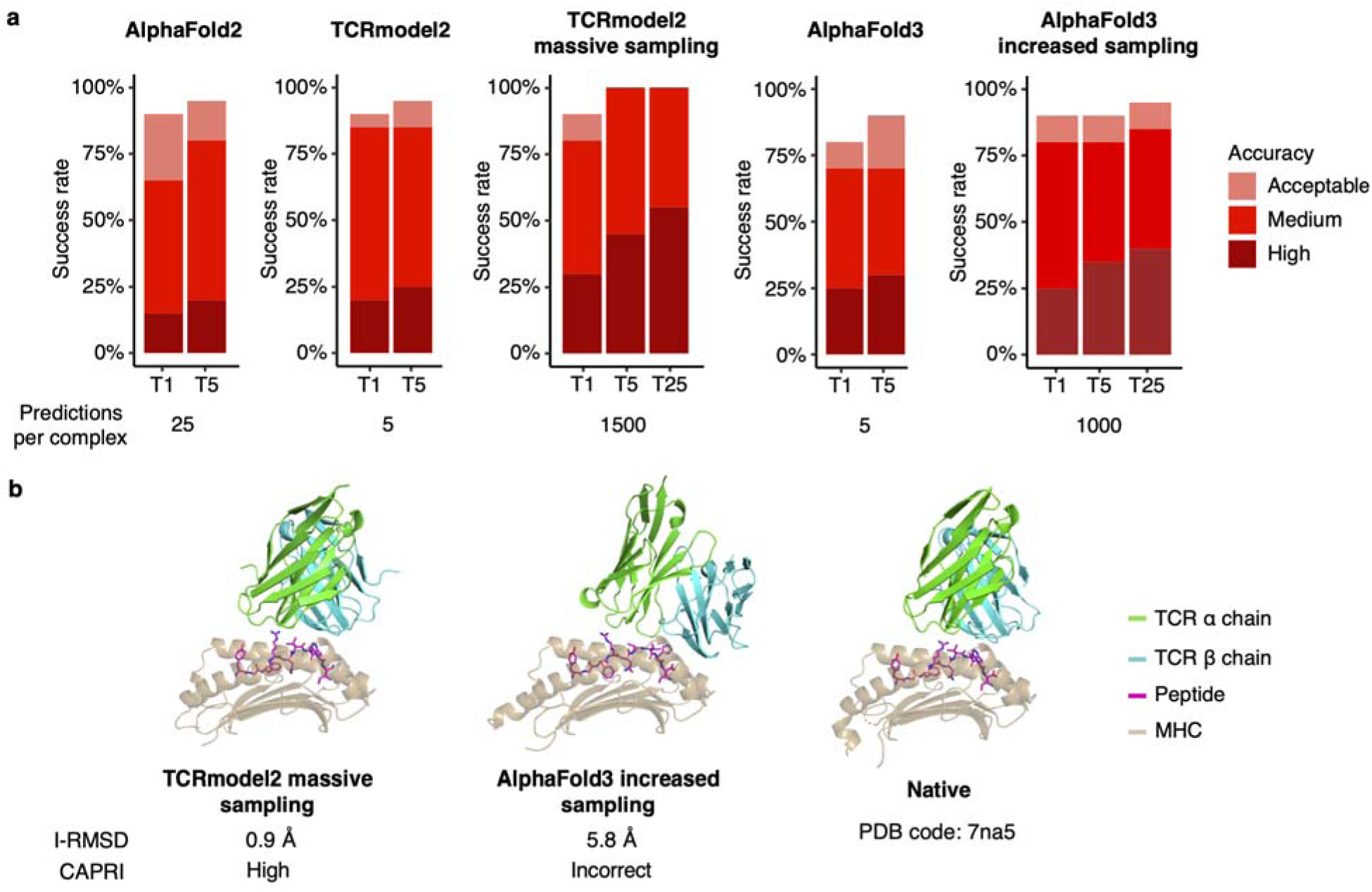
AlphaFold success for TCR-pMHC complex modeling. (a) Success rates for AlphaFold2, TCRmodel2, and AlphaFold3 with default and massive/increased sampling protocols for top-ranked (T1), top 5 (T5) and top 25 (T25, when available) ranked models per complex (20 test cases). Number of total models generated per complex are shown at bottom. Bars are colored by CAPRI accuracy levels, as shown on right. (b) Example top-ranked model of TCR-pMHC complex, with native complex structure (PDB code 7na5) on right. TCR, peptide, and MHC chains are colored as indicated on right. Interface RMSD (I-RMSD) from native complex and CAPRI accuracy level for each model are shown at bottom.

Given the observed improvements in model accuracy for top-ranked and highly ranked models across the immune recognition classes for increased sampling, we assessed the success rates for the full sets of models from each of those methods (T1000, T1500, or T8000; **Fig. S1**). This inclusion of all models showed that AF3 is able to generate high accuracy models for over 50% of complexes for antibody-protein and TCR-pMHC complexes. Those total success rates are notably higher or comparable to the AF2 antibody-protein and TCR-pMHC total success rates, respectively. Total near-native success rates (T1500 for AF2, T1000 for AF3) were lower for antibody-peptide complexes versus the other complex classes, as with the T1 success rates, indicative of higher modeling challenge for AF-based methods. While variable among AF versions and target complex classes, the differences between total and top-ranked success rates highlight opportunities for improved model ranking and selection methods to surpass the default AF2 and AF3 confidence-based ranking scores.

### Confidence scores and model accuracy

To investigate which AlphaFold2 and AlphaFold3 confidence scores are better predictors of antibody-protein, antibody-peptide, and TCR-pMHC model accuracy, we assessed and compared the classification accuracy of confidence scores for sets of pooled models from the massive and increased sampling benchmarking. Confidence scores compared were interface predicted TM score (ipTM), interface predicted LDDT score (I-pLDDT), and the AlphaFold-Multimer model confidence formulation, which is a linear combination of full complex predicted TM (pTM) and ipTM (0.8*ipTM + 0.2*pTM). We refer to the latter score as “model confidence score” in this study for brevity and in accordance with previous work^31^. Additionally, for AlphaFold3 and TCRmodel2 the receptor-ligand ipTM (RL-ipTM) score was assessed, as that score was readily available from the output of those algorithms. RL-ipTM score (alternatively referred to as Ab-Ag or TCR-pMHC ipTM) is the ipTM score for only the antibody-antigen and TCR-pMHC interfaces, rather than the ipTM between all chains present in the complex.

To assess the effective identification of near-native immune complex models, receiver operating characteristic area under the curve (ROC AUC) values for classification of high accuracy versus lower accuracy and incorrect models were compared (**Fig. 4, Table S4**). While differences in accuracy class proportions (**Table S4**) could potentially affect inter-protocol and inter-class comparisons, ROC AUC is viewed as generally robust to class imbalance scenarios ^50^. Our analysis revealed that all tested AF2 confidence-based scores had high near-native discrimination across immune receptor interfaces (AUC > 0.9), with strongest overall classification performance for antibody-protein interfaces (**Fig. 4A**). I-pLDDT showed higher classification performance versus the other metrics, particularly for antibody-peptide and TCR-pMHC interfaces. This may be partly expected due to the other inter-chain interfaces (i.e. antibody heavy-light chain for antibody-peptide complexes, peptide-MHC and TCRα-TCRβ chain interfaces for TCR-pMHC complexes) representing a sizable portion of overall ipTM and model confidence scores, whereas I-pLDDT is focused specifically on the respective receptor-ligand interface. For AF3, ROC AUC scores were high for antibody-protein interfaces (0.94 or above), as with AF2, however the classification accuracy dropped considerably for antibody-peptide and TCR-pMHC interfaces. Interface-specific AF3 metrics RL-ipTM and I-pLDDT performed better than model confidence and ipTM scores for TCR-pMHC interfaces, while RL-ipTM outperformed the other confidence scores for TCR-pMHC high accuracy model discrimination (ROC AUC = 0.82).

**Figure 4.**
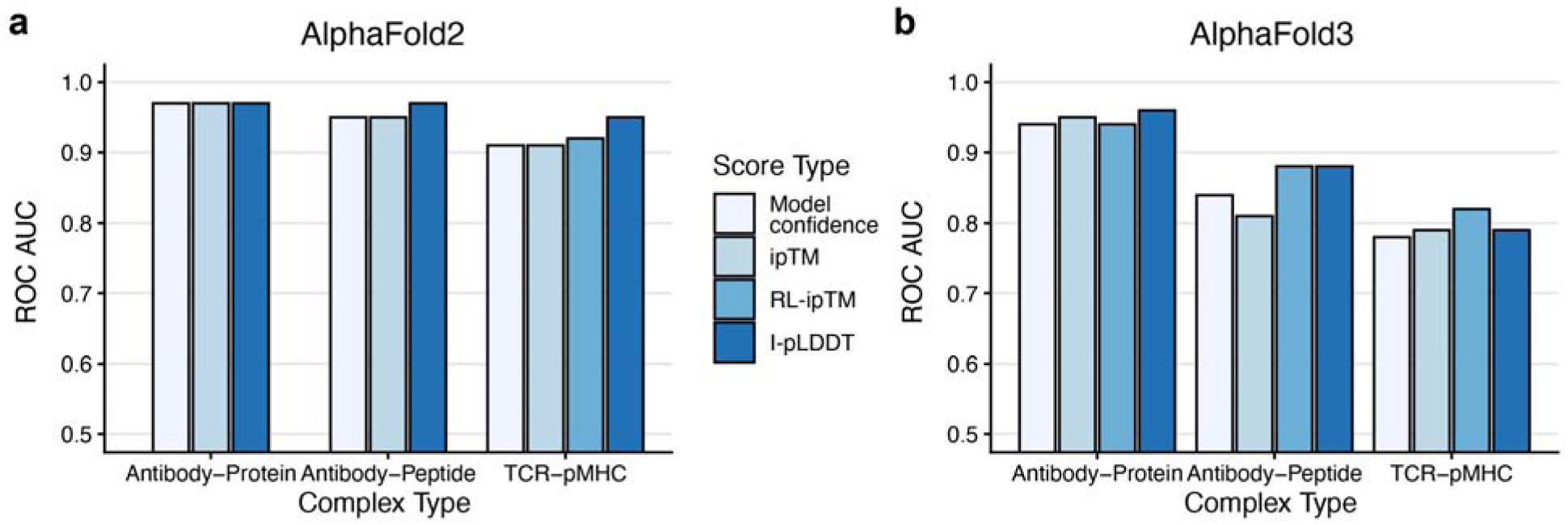
AlphaFold2 and AlphaFold3 confidence score model accuracy classification performance. Receiver operating characteristic area under the curve (ROC AUC) values for classification of high accuracy versus other models in pooled sets of AF2 and AF3 models from massive/increased sampling for antibody-protein, antibody-peptide, and TCR-pMHC complexes. Scores assessed were model confidence score (“Model conf.”), ipTM, receptor-ligand ipTM (“RL-ipTM”), and I-pLDDT. RL-ipTM score was not assessed for antibody-protein and antibody-peptide complexes in AF2 as that score is not natively output by AF2. TCRmodel2, which does output RL-ipTM score, was used for TCR-pMHC complex AF2 modeling in this analysis.

The distinction between AF2 and AF3 scoring is evident upon inspection of distributions of I-pLDDT and ipTM scores among model accuracy levels and interface classes (**Fig. 5**, **Fig. S2**). Particularly for antibody-peptide and TCR-pMHC interfaces, there is a considerable overlap between high and medium accuracy model scores for AF3, whereas there is less overlap for AF2 scores. However, AF3 confidence scores generally perform much better at discriminating medium and high accuracy models from lower accuracy models, in many cases outperforming AF2, as seen in the score distributions (**Fig. 5**, **Fig. S2**) as well as ROC AUC values (**Fig. S3**, **Table S5**). In accordance with the medium or higher accuracy classification ROC AUC values, AF3 confidence scores were found to have generally higher overall correlations with DockQ model accuracy values than AF2 confidence scores (**Table S5**).

**Figure 5.**
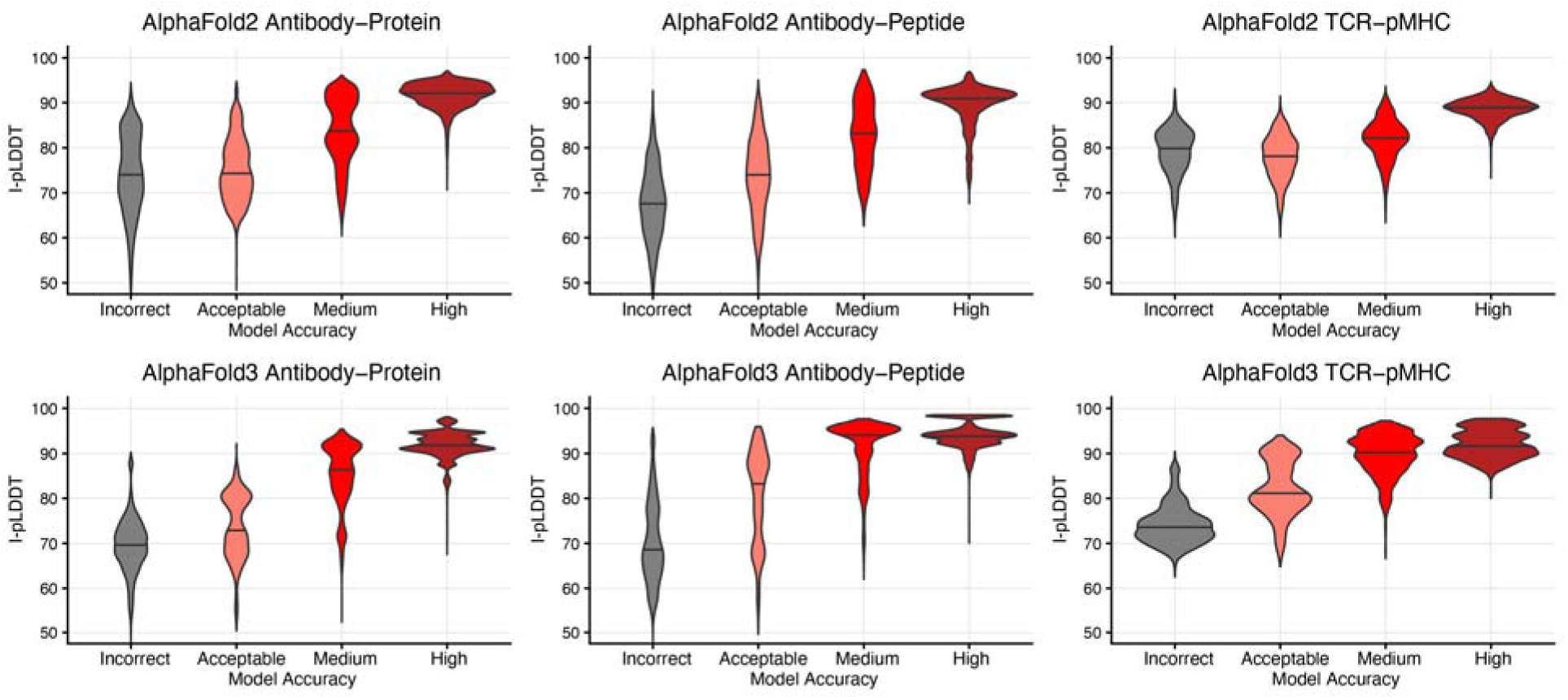
Distributions of I-pLDDT scores for model accuracy levels. AlphaFold2 (top) and AlphaFold3 (bottom) interface pLDDT (I-pLDDT) scores are shown for pooled sets of antibody-protein (left), antibody-peptide (middle), and TCR-pMHC models, classified by CAPRI model accuracy level.

To provide insights into confidence score cutoffs for selection of accurate models, we calculated success rates binned by score ranges for the three interface classes for I-pLDDT, ranking confidence, and RL-ipTM scores (**Fig. 6**, **Figs. S4-S7**). As seen in the I-pLDDT score binned success (**Fig. 6**), while success rates increase for higher I-pLDDT scores as expected, there are notable differences between AF2 and AF3, as well as the interface classes. For instance, AF3 I-pLDDT in the 90-92.5 score range has a greater than 50% rate of high accuracy models, whereas AF2 is near 25%, and still below 50% in the next I-pLDDT score bin (92.5-95). Conversely, both antibody-peptide and TCR-pMHC complex models in the 90-92.5 I-pLDDT score range have >50% high accuracy rates for AF2, while AF3 high accuracy rates are below 50% for that range and not surpassing 50% for higher score bins. Of note, the TCR-pMHC ranking confidence and RL-ipTM binned success distributions were less divergent for AF2 and AF3 (**Fig. S6**, **Fig. S7**). The RL-ipTM stratified success rates (**Fig. S7**) indicate that score cutoffs for that metric for all three interface classes in the 0.8-0.9 range would be sufficient for approximately 50% high accuracy model rates across the interface classes and AF protocols tested.

**Figure 6.**
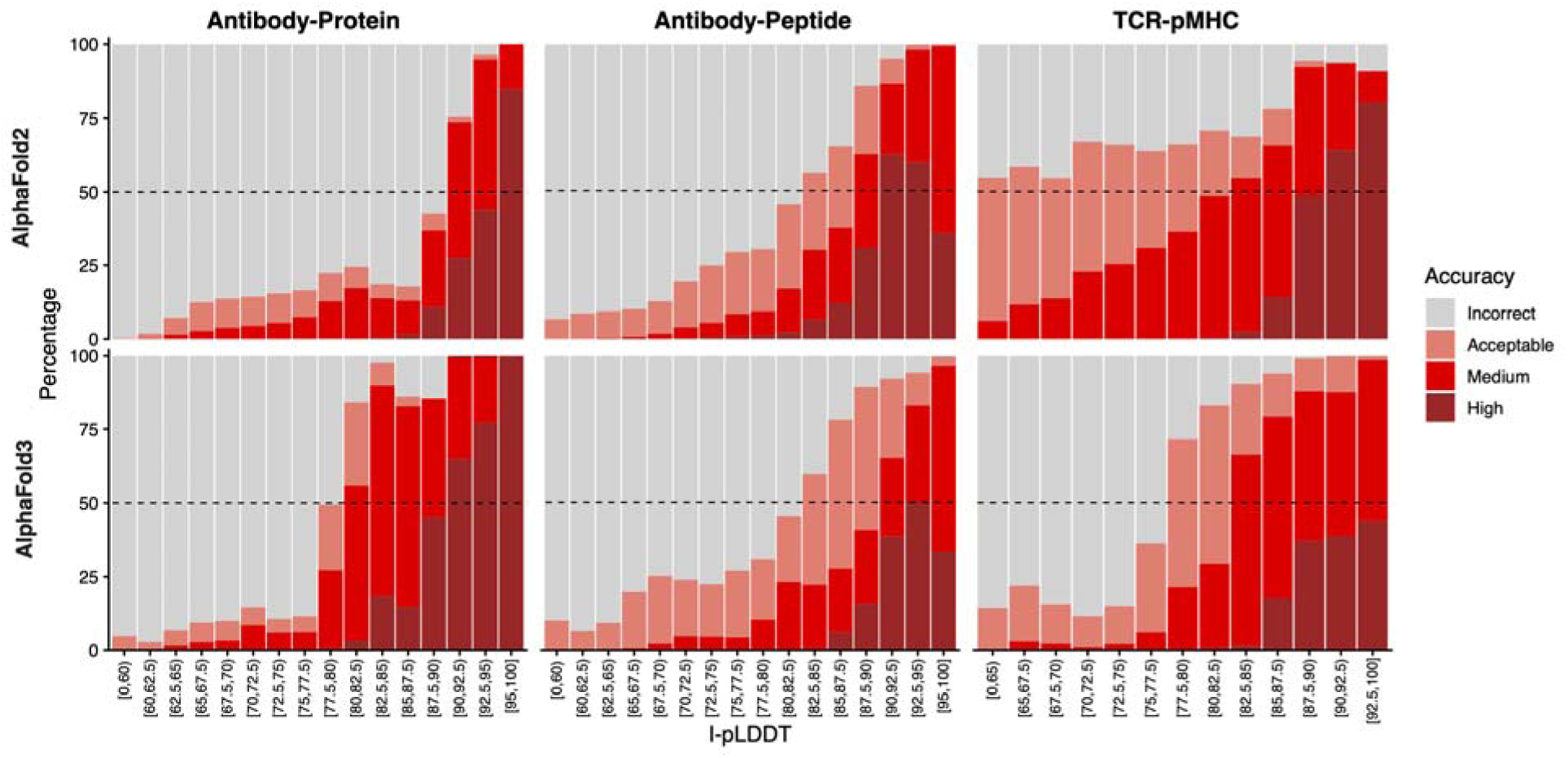
Model accuracy rates across I-pLDDT score ranges. AlphaFold2 (top) and AlphaFold3 (bottom) models were binned by I-pLDDT confidence scores for antibody-protein (left), antibody-peptide (middle), and TCR-pMHC (right) complexes, and percentages of CAPRI model accuracy levels in each score bin are indicated by bar colors, as shown on the right. Dashed lines at 50% percentage levels are shown for reference.

To analyze potential confidence score cutoffs through another metric, we calculated 50% high accuracy positive predictive value (PPV, or precision) score cutoffs and recall rates at those cutoffs (**Table S6**). Overall, the score cutoffs and recall rates varied considerably among interface classes and AF versions, which may be attributed to the different overall high accuracy model propensities in the pooled model sets (**Table S3**), for instance between AF2 and AF3 antibody-protein models (2.8% versus 19.8%, respectively). Additionally, uneven distributions of number of models among score ranges (**Figs. S4-S7**), including in ranges above the 50% PPV cutoff values, may complicate practical interpretation of some of the PPV-based cutoffs. Still, some score cutoffs seem more useful in predictive scenarios, such as the AF2 TCR-pMHC I-pLDDT cutoff of 87, which is quite similar to the 87.5 I-pLDDT score cutoff used in recent studies using AF2 (TCRmodel2) to model recently determined and structurally uncharacterized TCR-pMHC complexes^46, 47^.

### Comparison with other recent deep learning methods

Following the release of AlphaFold3, several open-source deep learning structure prediction methods were developed that are based on the reported AlphaFold3 architecture ^30^, including Boltz-1 ^40^ and Chai-1 ^39^. We tested both of those methods in comparison with AF3 for antibody-protein, antibody-peptide, and TCR-pMHC modeling accuracy, running each with 20 seeds per complex, corresponding to 100 ranked structures per complex (**Fig. 7**). This corresponds to moderate increased sampling, although fewer than the 200 seeds (1000 structures per complex) tested earlier in this study due to computational limitations. For TCR-pMHC modeling, we utilized an expanded set of 25 test cases versus our original set of 20 test cases (**Table S3**) and also included TCRmodel2 in the success comparison due to its comparable performance to AF3 (**Fig. 3**). The success comparison (**Fig. 7**) shows that AF3 generally outperforms Boltz-1 and Chai-1 for all three classes of interfaces. This is in agreement with previous comparative benchmarking of antibody-protein modeling ^51^, while demonstrating that AF3 is likewise superior to Boltz-1 and Chai-1 for antibody-peptide and TCR-pMHC complexes. For the latter classes of interfaces, Boltz-1 generally outperformed Chai-1, while for TCR-pMHC modeling, TCRmodel2 success was comparable to AF3, as seen previously with the moderately smaller benchmark with different seeds and sampling (**Fig. 3**).

**Figure 7.**
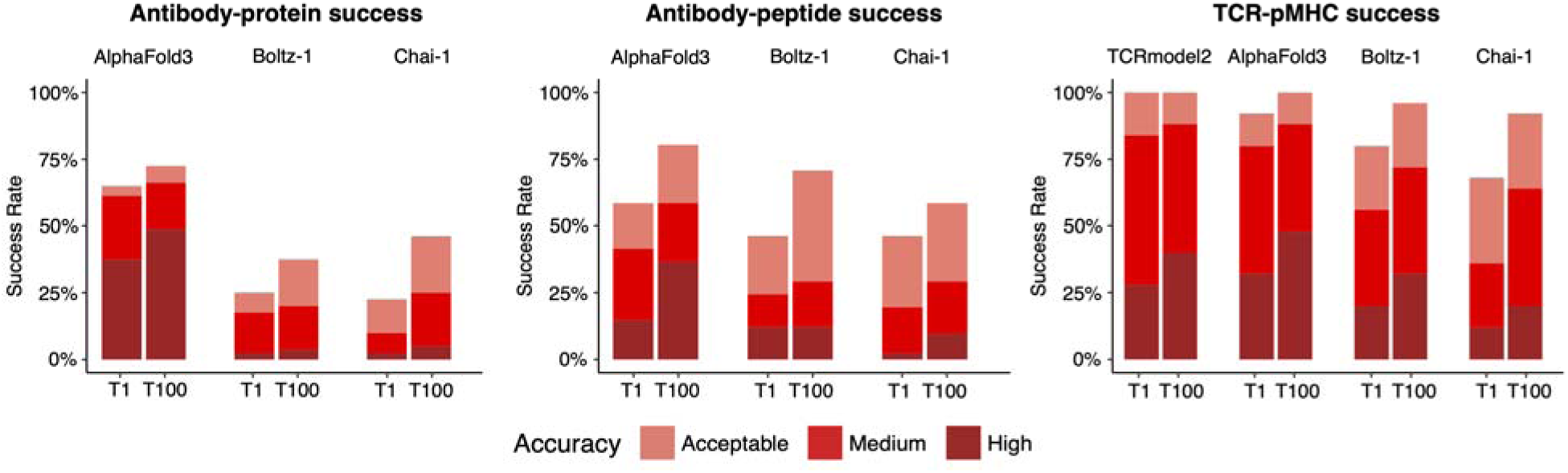
Comparison of AlphaFold3, Boltz-1, and Chai-1 success rates for complex classes. AlphaFold3, Boltz-1, and Chai-1 generated models for antibody-protein (left; N=80), antibody-peptide (middle; N=41), and TCR-pMHC (right; N=25) test cases, each producing 100 structures per complex (20 seeds), and success rates for top-ranked (T1) and full set of 100 models (T100) are shown, colored by accuracy as shown at bottom. Additionally, TCRmodel2 success is shown for the TCR-pMHC complexes, for comparison.

To assess additional recently released deep learning modeling tools, we performed a comparison of AF3, TCRmodel2, Boltz-1 with OpenFold3^52^, Protenix v1^53^, and SeedFold^54^, as well as the TCR-pMHC deep learning modeling tool tFold-TCR^55^ for modeling accuracy with the expanded TCR-pMHC benchmark set (**Fig. S8**). Modeling was performed with single seeds (1 or 5 structures generated per complex) to provide a balanced comparison, given that one method (tFold-TCR) only generates one structure per input. Comparison of top-ranked (T1) success from the methods showed that most had comparable rates of high accuracy models, except for AF3 which had a higher rate, and tFold-TCR that had no models generated with that accuracy. For medium or higher accuracy, TCRmodel2, AF3, and SeedFold demonstrated the highest rates of T1 success (∼75%) versus the other methods.

### Complementarity between deep learning methods

An intriguing possibility for immune complex modeling is the potential complementarity between methods, which would enable improvement in modeling success rates versus any single top-performing protocol through the use of pooled models from different deep learning protocols. To investigate this, we analyzed the per-complex top-ranked model accuracies for TCR-pMHC (**Fig. 8a**), antibody-peptide (**Fig. 8b**), and antibody-protein (**Fig. S9**) complexes from the above AF3, Boltz-1, and Chai-1 results. TCRmodel2 (TCR-pMHC) and AF2.3 (antibody-peptide, antibody-protein) results with 20 seeds are also included in the comparisons, as AF2-representative protocols. Comparing per-model accuracies revealed notable complementarity between methods, particularly between AF2 and AF3, with limited additional contributions from the AF3-related methods Boltz-1 and Chai-1. For TCR-pMHC complexes, combining TCRmodel2 and AF3 results in high accuracy models for 11 out of 25 complexes (44%), an improvement over the top-performing single method (AF3; 8 out of 25 complexes, 32%). High accuracy complexes increase by 8 cases, from 30% (AF3 alone) to 38% (all methods) for antibody-protein complexes, while greater relative improvement was observed for antibody-peptide complexes, from 6/41 cases (15%: AF3 alone) to 13/41 cases (32%; all methods). Complementarity between deep learning methods for near-native (medium or higher) accuracy was highest for antibody-peptide complexes, which increased from 17 (41%; AF3) to 24 (59%) cases when considering all methods. Near-native success increased by 8 cases (46% to 56%) and by 3 cases (84% to 96%) for antibody-protein and TCR-pMHC cases, respectively, for pooling versus the best individual methods (AF3, TCRmodel2). This complementarity highlights an opportunity to increase success through pooling of top or highly ranked models from AF2, AF3, and other protocols, while also showing a remaining set of targets in each class with no top-ranked high or near-native accuracy model from any method.

**Figure 8.**
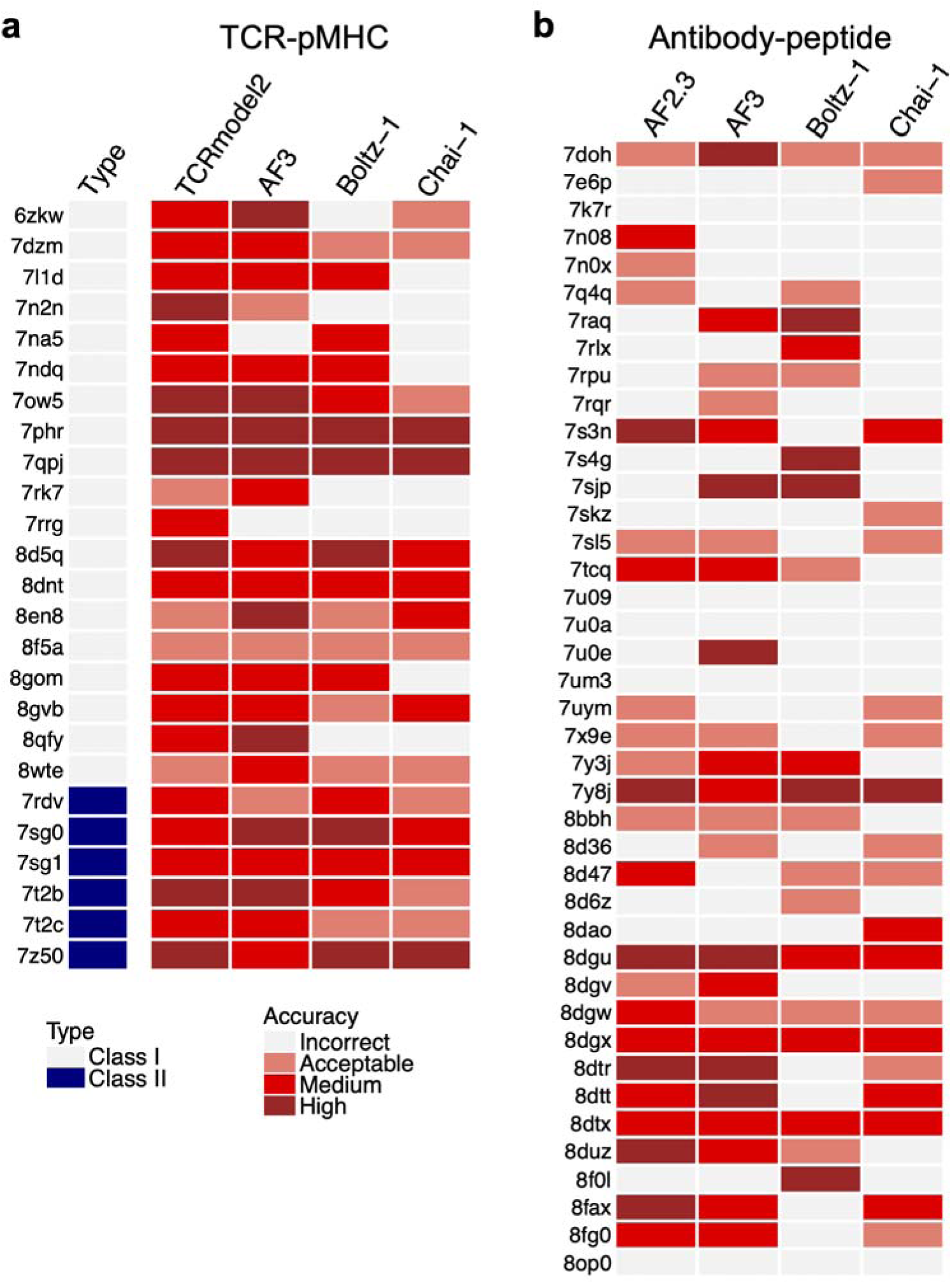
Top-ranked model per-case success for TCR-pMHC and antibody-peptide complexes. Top-ranked model accuracy for individual (a) TCR-pMHC and (b) antibody-peptide test cases for AF3, Boltz-1, Chai-1, as well as TCRmodel2 (TCR-pMHC) or AF2.3 (antibody-peptide) is shown, with model accuracy levels colored as shown on bottom in (a). All methods generated 100 structures per complex, which were ranked by model confidence score to obtain the top-ranked model. Test cases are denoted by complex PDB code (left), and in (a) Type denotes the MHC class of the TCR-pMHC complex.

### Modeling a noncanonical TCR-pMHC complex

To test the capability of AlphaFold and related deep learning methods to model an immune recognition scenario far outside of the training set, we modeled the structure of a human TCR in complex with Class II MHC and peptide that represents a highly unusual binding mode which differs markedly from other structurally characterized TCR-pMHC interactions, with the TCR engaging the side of the MHC rather than the face with the peptide and both MHC helices^56^ (**Fig. 9A**). We performed modeling of that complex with multiple deep learning methods (AF3, TCRmodel2, Boltz-1, OpenFold3, and SeedFold) generating 100 models per complex, assessing model accuracy versus the recently reported X-ray structure (PDB code 9EJG)^56^. Strikingly, only AF3 generated any accurate models, producing multiple high accuracy models within its set of 100, including one ranked third (**Fig. 9B**, **Table S7**); no other method produced an accurate model at any rank. The third-ranked AF3 model had 0.97 Å I-RMSD from the X-ray structure, and high AF3 confidence scores (0.876 model confidence, 93.1 I-pLDDT), while the top-ranked AF3 model based on AFM ranking score had an incorrect binding mode reflective of a canonical TCR-pMHC interaction (**Fig. 9C**), with slightly higher model confidence score (0.892), but lower I-pLDDT score (89.9) versus the high accuracy AF3 model. The other tested methods generated inaccurate models with canonical-like binding modes as with the top-ranked AF3 model, or non-canonical erroneous binding modes (**Fig. 9D**). Of note, the top-ranked models from the other approaches had lower model confidence and I-pLDDT scores than the high accuracy AF3 third-ranked model (**Table S7**), and that AF3 model had the highest I-pLDDT score out of the set of 100 AF3 models. This highlights the potential of interface-focused scores such as I-pLDDT as well as model pooling, although use of I-pLDDT score alone has limitations based on previous antibody-protein modeling^37^. Overall, while deep learning structure prediction methods can accurately model unseen complexes that are highly distant from the training set, performance is variable, and model selection and ranking requires careful consideration and improvement.

**Figure 9.**
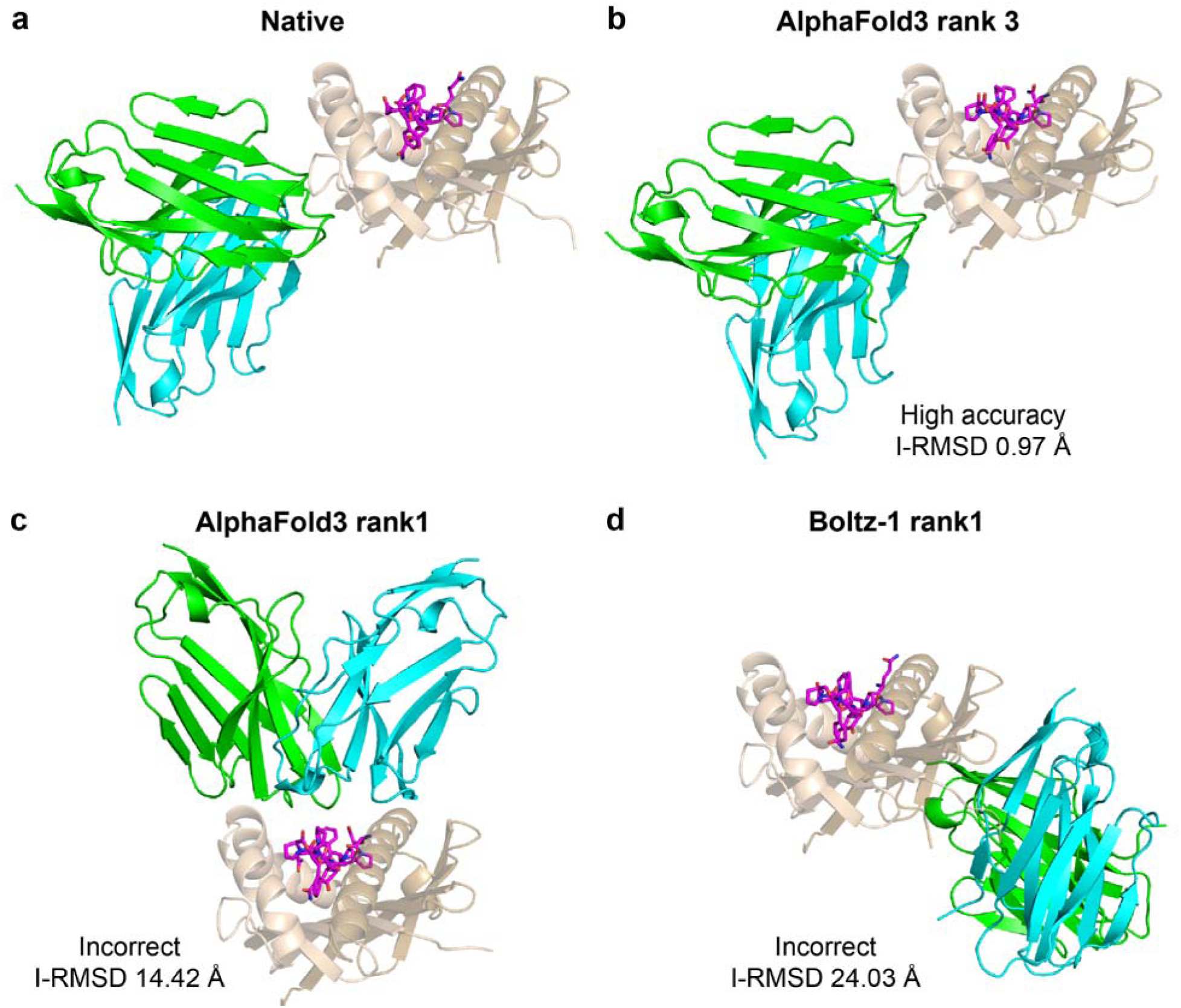
Modeling of a noncanonical TCR-pMHC complex. (a) X-ray structure of the G9 TCR in complex with the HLA-DQ2.5 and peptide, showing a noncanonical peptide independent TCR-pMHC binding mode (PDB code 9EJG)^56^, (b) a high accuracy structural model from AlphaFold3 (rank 3), (c) an incorrect AlphaFold3 (rank 1) model with canonical TCR-pMHC binding mode, (d) an incorrect Boltz-1 (rank 1) model of the complex. Structures are colored by chain, with TCR α chain green, TCR β chain cyan, MHC α dark tan, MHC β light tan, and peptide magenta. CAPRI accuracy level and I-RMSD from the native complex structure are shown below each model in (b-d).

## Discussion

In this study we performed extensive comparative benchmarking of multiple deep learning methods and protocols on three distinct classes of immune recognition, identifying optimal protocols, differential success rates, and highlighting considerations for scoring and model selection. While AF3 was found to outperform other AF3-like methods and AF2 for antibody-protein complexes, in accordance with what others previously observed ^38, 51, 57^, its advantage over AF2 was less pronounced or nonexistent for TCR-pMHC and antibody-peptide interaction modeling. The lowest overall modeling success rates were observed for antibody-peptide complexes, highlighting the ongoing challenge presented by that key class of interactions, while antibody-protein and TCR-pMHC complexes still contained substantial fractions of complexes without high accuracy models generated.

Given the differential modeling accuracy between complexes of a given class, a major question is what determinants or features underlie successful versus unsuccessful deep learning structural modeling. Previously we found some association of antibody type (nanobody versus heavy-light chain antibody), glycosylation near binding interface, and computed interface energetic score with modeling success for antibody-protein interactions^31^. Others have noted that overlapping structural features between training and test sets is associated with antibody-protein modeling success for AlphaFold2^58^ as well as AlphaFold3^57^, and a similar possible bias was also found for protein-peptide complex modeling in AlphaFold2 and AlphaFold3^59^. Even with these limitations, AlphaFold appears to have learned some immune recognition principles and is clearly capable of modeling previously unseen complexes in some cases, as seen in sizable fractions of benchmark complexes in this study, as well as previous anecdotal modeling of newly determined antibody-protein complexes^60^, TCR-pMHC complexes^46, 47^, and an antibody-peptide complex^37^, and exemplified by our modeling of the noncanonical TCR-pMHC complex in this study.

There are numerous opportunities for future developments to improve modeling success within or across the immune recognition classes. Improved model scoring and ranking is one major area of interest, given the difference between top-ranked and full set success observed in our benchmarking, as well as the potential benefit of pooled sets of models from multiple protocols also noted in this study. Indeed, the AF2 massive sampling approach represents pooled structures from multiple AF2 models (v.2.1, 2.2, and 2.3), and the developers of one implementation noted the need for improved pooled set scoring versus default AF2 confidence scores for antibody-antigen modeling ^36^. Several adaptations of AlphaFold2 and AlphaFold3 confidence scores have been generated to improve accuracy or interaction prediction for modeled complexes^61–63^, while two recently developed antibody-antigen deep learning-based scoring protocols, DeepRank-Ab^64^ and ABAG-Rank^65^, reportedly outperform AF3 for antibody-antigen model selection. Additionally, modifications to AF2 and AF3 to score previously generated sets of structures ^66, 67^ could potentially be used or adapted to score pooled sets of complex models from different protocols. Others have developed hybrid pipelines that utilize AlphaFold in conjunction with physics-based approaches^68^, showing utility in a protein docking benchmark set^69^ and predictive complex modeling in a recent CASP/CAPRI round^70^. Finally, it is possible that a more generalizable model could be trained, for instance through fine-tuning the AF3 model using an open-source framework such as Boltz-1^40^ or OpenFold3^52, 71^, although the limits in available PDB structures would require immune complex distillation strategies (for instance using high confidence modeled structures), as employed in AF3 training with AF2-modeled monomer structures^30^, and recently explored by the Boltz team with TCR-pMHC complexes^72^. Recently reported deep learning algorithms Protenix-v2^73^, IsoDDE^74^, and ESMFold2^75^ have reported success rates higher than AF3 for antibody-protein complex modeling based on the authors’ benchmarking, although independent benchmarking would be needed to confirm that, and one of those methods is not currently available to the public.

One intriguing possibility based on the increasing success of deep learning protocols is the prediction of immune receptor targets from sequence. This key challenge has been noted by others, particularly regarding the prediction of TCR specificity^76, 77^, as state-of-the-art methods struggle with “unseen” epitopes not present in training^78, 79^. A recent TCR specificity prediction challenge with unseen epitopes, IMMREP25, showed that several teams and approaches that used AF3 models and confidence scores of candidate TCR-pMHC complexes were able to predict specificities at higher rates than random background and other approaches^80^. Those successes highlight the promise of AF3 and possible future versions or adaptations to more accurately and generally predict TCR specificities from the sequence of the receptor alone, enabling de-orphanization and specificity assignment of TCRs of interest (versus experimental approaches^81, 82^) and insights into disease and TCR-based biotherapeutics.

There are some limitations to this study. Due to the availability of structures in the PDB, the sizes of the antibody-peptide (41 cases) and TCR-pMHC (20 or 25 cases) benchmark sets are more limited than for the antibody-protein set (80 cases). Additionally, computational limitations, and algorithm availability only as a web server in some cases, prevented testing of greater numbers of seeds and sampling for non-AlphaFold methods.

In summary, our benchmarking identifies increased sampling in AF2 and AF3, along with the use of interface-focused confidence scores such as I-pLDDT and RL-ipTM as ways to optimize antibody-protein, antibody-peptide, and TCR-pMHC modeling success, while pooling models from different deep learning protocols can provide further improvements. However, all tested classes of complexes showed some challenge, most notably antibody-peptide complexes for which approximately half of the complexes did not have near-native top-ranked models generated. Even with these limitations, the accuracy of current approaches and model confidence predictions show increasing utility of deep learning methods in predictive immune complex modeling scenarios, while highlighting the need for future developments. Such advances will provide the opportunity to more generally and accurately gain insights into antibody and TCR recognition, predict specificities, and computationally design and screen targeted antibody- and TCR-based immunotherapeutics de novo^83–87^.

## Methods

### Benchmark assembly

A nonredundant set of antibody-protein benchmark test cases was assembled from antibody-protein PDB structures obtained from the SAbDab database^33^. Structural filtering and nonredundancy criteria were based on previous antibody-protein benchmark assembly^31^. Specifically, the criteria were: 1) Structural resolution 3.0 Å or better, 2) Nonredundant with each other, and with antibody-antigen complexes with resolution ≤ 9 Å and release date from September 30, 2021 or before. Nonredundancy is defined as heavy chain V domain sequence identity < 90% and full variable domain sequence identity < 90%. Alternatively, complexes are considered nonredundant with no sequence match between antigens using BLAST^88^ with default parameters. Complexes passing these filters were additionally analyzed for structural redundancy, in which pairs of structures with < 5 Å heavy chain Cα atom RMSD after superposition of antigens using FAST structural alignment^89^, along with >70% identity between heavy chain V domain, light chain V domain, or concatenated CDR loop sequences, were considered redundant. Finally, manual structural inspection of remaining cases was performed to remove noncanonical complexes (e.g. with constant domain binding to an antigen), complexes with large or interface-proximal unresolved regions, and complexes with HETATMs forming a key part of the antibody-antigen interface.

The antibody-peptide complex benchmark set was assembled with available antibody-peptide structures in the PDB, identified through the SAbDab database. Structures were filtered with a protocol that is similar to what was used previously for antibody-peptide structure dataset curation ^41^. Specifically, criteria for filtering were: 1) Structural resolution of 3.0 Å or better, 2) No internal unresolved residues in the peptide, 3) Resolved peptide length not less than 5 residues and not greater than 25 residues, 4) No non-protein (HETATM) residues or atoms directly in the antibody-peptide interface, based on automated analysis followed by manual inspection, 5) Not redundant with other complexes within the dataset, and with any previously released antibody-antigen (protein or peptide antigen) complexes that have a structural resolution of 9.0 Å or better and release date on or before the September 30, 2021 cutoff. Nonredundancy is defined as having less than 90% identity in the sequence of the heavy chain variable domain as well as the concatenated variable domain sequences.

The TCR-pMHC complex benchmark set of 20 cases was assembled previously^28^. Complexes were identified from the PDB in the TCR3d database^34^ based on the following criteria: 1) Structure resolution of 3.25 Å or better and 2) No redundancy with any TCR-pMHC complexes within the set and with TCR complexes released on or before the September 30, 2021 date cutoff. Nonredundancy is defined as having < 95% sequence identity in the TCR α and β chain variable domains and full variable domain sequence identity < 92% with respect to any TCR-pMHC complex of the same class. The updated set of TCR-pMHC complexes includes five additional complexes which were identified from available TCR3d and PDB structures in July 2024.

All complex structures were modeled and assessed using antibody or TCR variable domains (no constant domains included), for consistency across complexes, computational efficiency, and following previous benchmarking practices^28, 31^. For TCR-pMHC complexes, only the peptide-binding domains of the MHC proteins (α1α2 for Class I, α1β1 for Class II) were included in modeling and assessment. For Class II TCR-pMHC complexes, the peptide 9-mer core plus one residue on each terminus was used (11 residues total), as in previous work^28^.

### AlphaFold2

To generate default AlphaFold v2.3 predictions, we utilized AlphaFold v2.3 was downloaded from its Github repository (https://github.com/google-deepmind/alphafold) in February 2023 and installed on a local computer cluster. A template date cutoff of September 30, 2021 was used, and all other settings were default, giving 25 predictions generated per complex.

AlphaFold2 massive sampling for antibody-protein complexes was performed using an adaptation of the AFsample algorithm^90^. We utilized the AFsample protocol and code (https://github.com/bjornwallner/alphafoldv2.2.0), making modifications to include AlphaFold v2.3 models in the pooled set as described previously. 8000 predictions were generated per complex using AlphaFold v2.1, v2.2, and v2.3:

- 1000 predictions with v2.1, with templates and full dropout
- 1000 predictions with v2.2, with templates and full dropout
- 1000 predictions with v2.1, without templates and with Evoformer dropout
- 1000 predictions with v2.2, without templates and with Evoformer dropout
- 1000 predictions with v2.1, without templates and with Evoformer dropout, 21 recycles
- 1000 predictions with v2.2, without templates and with Evoformer dropout, 9 recycles
- 1000 predictions with v2.3, with templates and full dropout
- 1000 predictions with v2.3, without templates and with Evoformer dropout

Where not specified above, the number of recycles was default (3 recycles for v2.1, 2.2, and 20 recycles with early stopping for v2.3). Following the AFsample protocol, we performed Amber relaxation on the top 5 predictions, ranked by model confidence.

For massive sampling modeling of antibody-peptide complexes, 1500 predictions were generated per complex. 500 predictions each from AlphaFold v2.1, v2.2, and v2.3 were generated, with no templates, and Evoformer dropout.

Default AlphaFold2 and pooled massive sampling models were ranked based on the default AlphaFold2 ranking score, AlphaFold-Multimer model confidence score (0.8*ipTM + 0.2*pTM), with top-ranked predictions Amber relaxed based on default behavior. For all protocols in which templates were enabled, the template date cutoff was set to September 30, 2021.

### TCRmodel2

TCRmodel2 is a previously developed adaptation of AlphaFold2 to model TCR-pMHC complexes^28^. Default TCRmodel2 generates 5 structures per complex with AlphaFold v2.3 parameters. For TCRmodel2 massive sampling, 1500 structures per complex were generated, using 500 structures each from AlphaFold v2.1, v2.2, and v2.3 with Evoformer dropout and no TCR structural templates. Peptide-MHC templates, which are used by TCRmodel2 to enforce peptide geometry in the MHC groove, were retained for massive sampling. Peptide-MHC templates and TCR templates (default TCRmodel2 only) were used with a September 30, 2021 release date cutoff. As with AlphaFold2, all TCRmodel2 models were ranked using AF-Multimer model scores.

### AlphaFold3

To generate default AlphaFold3 predictions, we utilized the public web server (https://alphafoldserver.com/) in May 2024. One seed per complex was used, generating 5 predictions per complex. Seeds were generated at random, and the server only allowed chain templates from September 30, 2021 or before.

To generate larger sets of AlphaFold3 predictions per complex (increased sampling), we downloaded AlphaFold3 code from its Github repository (https://github.com/google-deepmind/alphafold3) in December 2024 and installed it locally. AlphaFold3 was run with 20 seeds (100 structures per complex) or 200 seeds (1000 structures per complex), with a template date cutoff of September 30, 2021.

We ranked AlphaFold3 models using the model confidence score (0.8*ipTM + 0.2*pTM), for consistency with the AlphaFold2 and TCRmodel2 confidence score-based model ranking. This is similar but not identical to the AlphaFold3 ranking score, which includes the model confidence score ipTM and pTM terms and weights, with two additional terms to favor disorder and disfavor clashes in models^30^.

### Other deep learning modeling methods

For other tested deep learning methods, the respective tools were downloaded from Github repositories and run locally. We downloaded Boltz-1^40^ (https://github.com/jwohlwend/boltz) in December 2024, Chai-1^39^ (https://github.com/chaidiscovery/chai-lab) in January 2025, OpenFold3-preview^52, 71^ (https://github.com/aqlaboratory/openfold-3) in November 2025, Protenix v1^53^ (https://github.com/bytedance/Protenix) in March 2026, and tFold-TCR^55^ (https://github.com/TencentAI4S/tfold) in March 2026. To generate Protenix v1 predictions, we used the protenix_base_default_v1.0.0 model, which has the same training date cutoff as AF3. To generate SeedFold^54^ predictions, we utilized its public web server (https://seedfold.io/dashboard/proteinPrediction; accessed March 2026), selecting the SeedFold_v1.0.0 model based on superior reported performance on antibody-protein complex modeling from its authors^54^.

Boltz-1, Chai-1, SeedFold, and tFold-TCR did not use structural templates, while OpenFold3 and Protenix-v1 were run with a September 30, 2021 template date cutoff. All methods generated five structures per seed for each complex by default, with the exception of tFold-TCR and Boltz-1 which generate one structure per complex by default. To generate five structures per seed per complex in Boltz-1, we changed its *diffusion_samples* parameter from one to five. To generate 100 models per complex for all methods, 20 random seeds were used, except tFold-TCR which does not enable random seeds. For consistency with AF2 and AF3 model ranking, models generated by each method were ranked by model confidence score (0.8*ipTM + 0.2*pTM).

### Model accuracy assessment

The accuracy of antibody-antigen and TCR-pMHC complex models was evaluated with DockQ v1.0^91^, downloaded from its Github repository (https://github.com/bjornwallner/DockQ). DockQ calculates the accuracy of models by comparing them to experimentally determined structures. It calculates interface and ligand backbone RMSD (I-RMSD and L-RMSD, respectively), the fraction of native contacts (fnat), and the DockQ score, which is a continuous metric (0-1) of complex model accuracy based on I-RMSD, L-RMSD, and fnat. Additionally, based on the model’s similarity to the native structure, the DockQ program determines the model’s CAPRI (Critical Assessment of Prediction of Interactions) accuracy, categorizing models into four distinct accuracy classes: “High”, “Medium”, “Acceptable” and “Incorrect” based on a combination of I-RMSD, L-RMSD, and fnat^49^. For assessment of antibody-peptide complex accuracy, the “-capri_peptide” command line option was used in DockQ, to utilize the CAPRI criteria for protein-peptide interfaces^49^.

### AlphaFold Interface pLDDT and RL-ipTM score calculations

Interface pLDDT (I-pLDDT) score was determined by computing the average pLDDT value for residues within a 4.0 Å distance cutoff at the antibody-antigen, antibody-peptide, or TCR-pMHC interface. For models lacking any residues within a 4.0 Å distance of the interacting partner, we assigned an I-pLDDT score of −1. For AlphaFold3 models, which have separate pLDDT values for each atom, residue pLDDT values were obtained by averaging atom pLDDT values for each residue.

Receptor-ligand ipTM (RL-ipTM) scores, reflecting the ipTM evaluated across the interface between receptor (TCR or antibody) and ligand (pMHC or antigen), were obtained for TCR-pMHC complexes from TCRmodel2 and for both TCR-pMHC and antibody-antigen complexes from AlphaFold3. To calculate the TCR-pMHC ipTM score in TCRmodel2, AlphaFold2 was modified to treat the TCR and pMHC within the predicted complex as a single chain each at the time of the ipTM score calculation, and that score is output by TCRmodel2^28^. Antibody-antigen RL-ipTM was not computed for AlphaFold2 models, as it is not readily obtained from the standard output. For AlphaFold3 models, RL-ipTM scores were calculated by averaging the chain_pair_iptm scores between antibody and antigen chains, or TCR and pMHC chains.

### Plotting and statistical analysis

Pymol (Schrödinger,Inc.) was used to generate structural figures. The R program (r-project.org) was used to generate all other figures, with the ComplexHeatmap R package^92^ used for heatmaps. Precision, recall, correlations, and ROC AUC values were calculated in Python using the scikit-learn and SciPy libraries.

## Acknowledgements

This work was supported by National Institutes of Health grant R35GM144083 (to B.G.P.), as well as seed grant funding through the National Cancer Institute-University of Maryland Partnership (to R.Y. and B.G.P.). The authors acknowledge the University of Maryland supercomputing resources (http://hpcc.umd.edu) made available for conducting the research reported in this paper, as well as the Institute for Bioscience and Biotechnology Research computing resources and IT staff. We also thank John Jumper (DeepMind) for information regarding the AlphaFold3 server and its template usage, as well as the SeedFold team for information on their server’s template usage. We are grateful to members of the Pierce laboratory, as well as Dr. Roy Mariuzza and Dr. Helder Ribeiro-Filho, for helpful comments and discussions.

## Data and code availability

Code for AlphaFold2, AlphaFold3, TCRmodel2, and other tested modeling tools are available as code and/or web server from their respective developers. Analysis scripts used in this study, and individual structural model accuracy data and coordinates, are available from the authors upon request. Benchmark test case lists are available as supplemental data (**Tables S1-S3**).

## Notes

### Competing Interest Statement

The authors have declared no competing interest.

