## Supplemental Files for "Benchmarking AlphaFold and related deep learning approaches for modeling antibody and TCR antigen recognition"

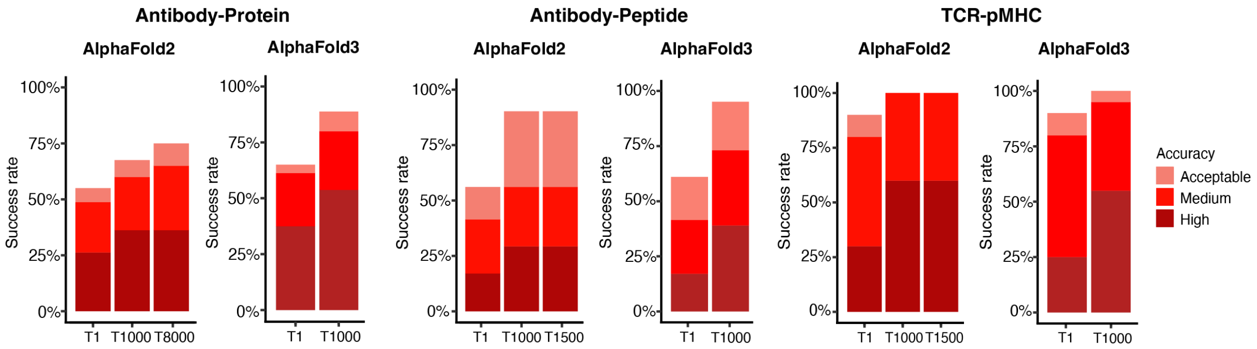


**Figure S1. AlphaFold top-ranked and total success for immune complex modeling.** Success rates for AlphaFold2 and AlphaFold3 with massive/increased sampling protocols for top-ranked (T1), top 1000 (T1000) models, and T1500/T8000 success for approaches in which 1500 or 8000 total models were generated, for antibody-protein (left), antibody-peptide (middle), and TCR-pMHC (right) complexes. For TCR-pMHC modeling, the TCRmodel2 adaptation of AlphaFold2 was used instead of default AlphaFold2. Bars are colored by CAPRI accuracy levels, as shown on right.

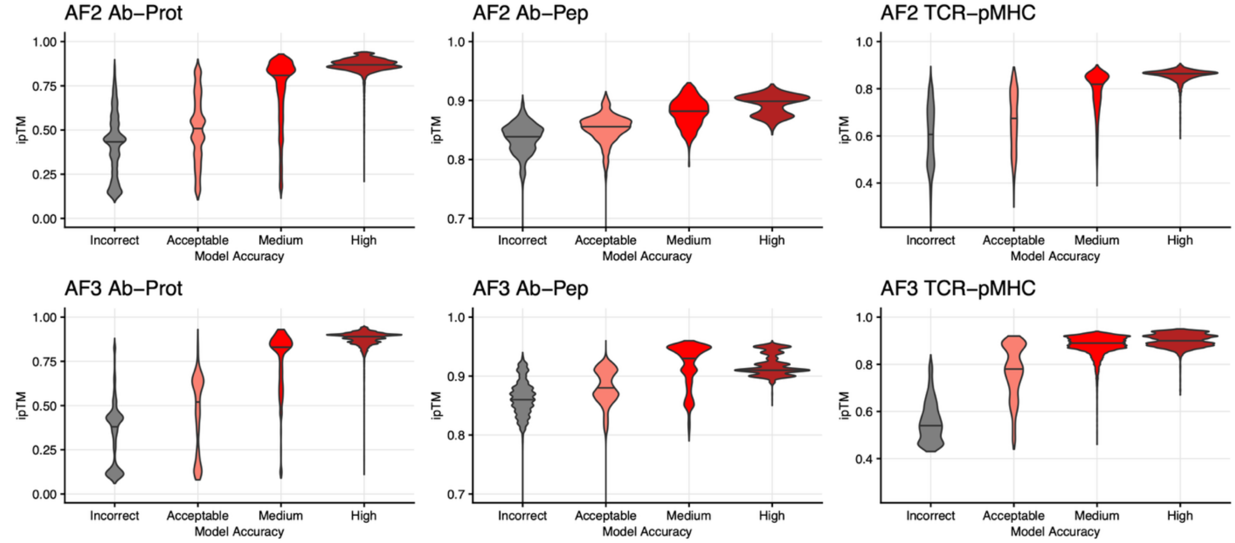


**Figure S2. ipTM score distributions across accuracy levels.** AlphaFold2 (top) and AlphaFold3 (bottom) ipTM scores are shown for pooled sets of antibody-protein (left), antibody-peptide (middle), and TCR-pMHC models, classified by CAPRI model accuracy level.

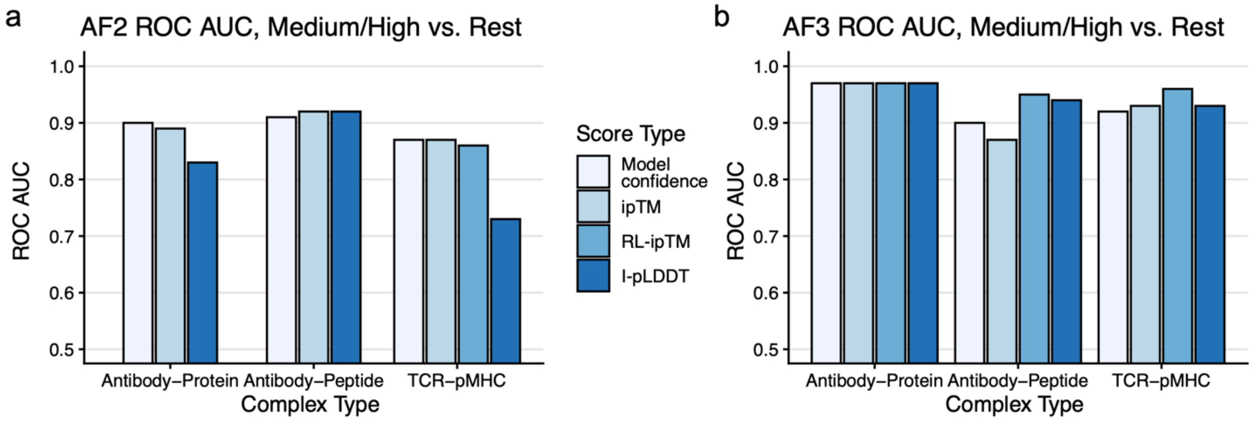


**Figure S3. AlphaFold2 and AlphaFold3 confidence score model accuracy classification performance for near-native versus other models.** Receiver operating characteristic area under the curve (ROC AUC) values for classification of near-native (medium or high CAPRI accuracy) versus other models (acceptable or incorrect CAPRI accuracy) in pooled sets of (a) AF2 and (b) AF3 models from massive/increased sampling for antibody-protein, antibody-peptide, and TCR-pMHC complexes. Scores assessed were model confidence score, ipTM, receptor-ligand ipTM ("RL-ipTM"), and I-pLDDT. RL-ipTM score was not assessed for antibody-protein and antibody-peptide complexes in AF2 as that score is not natively output by AF2. TCRmodel2, which does output RL-ipTM score, was used for TCR-pMHC complex AF2 modeling in this analysis.

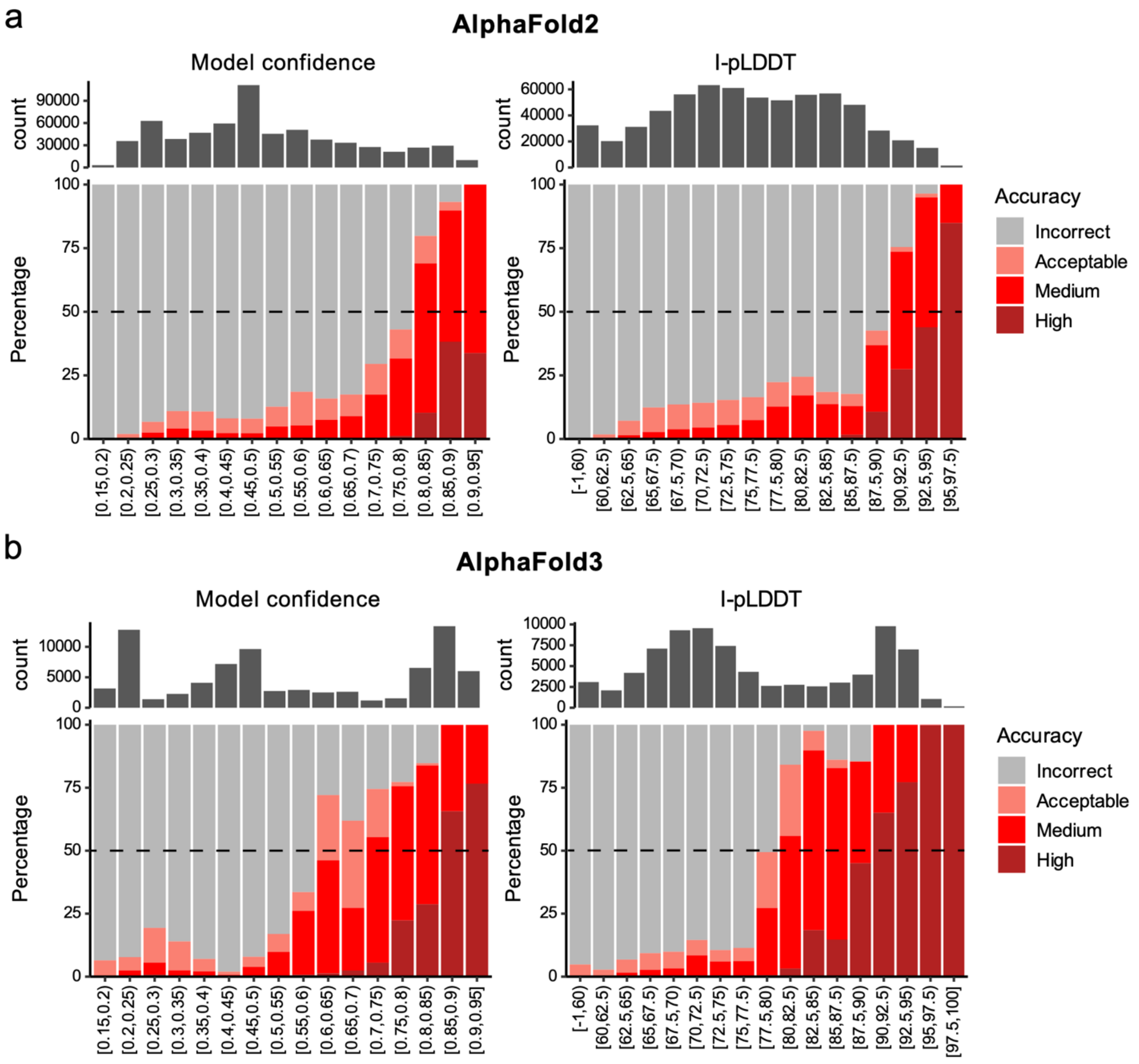


**Figure S4. Antibody-protein success rates for model confidence and I-pLDDT score ranges.** (A) AlphaFold2 and (B) AlphaFold3 success rates for pooled antibody-protein benchmark models are shown for model confidence (left) and I-pLDDT (right) score ranges, colored by CAPRI accuracy level as shown on right. Numbers of models per bin are shown at top, and dashed line at 50% is shown for reference.

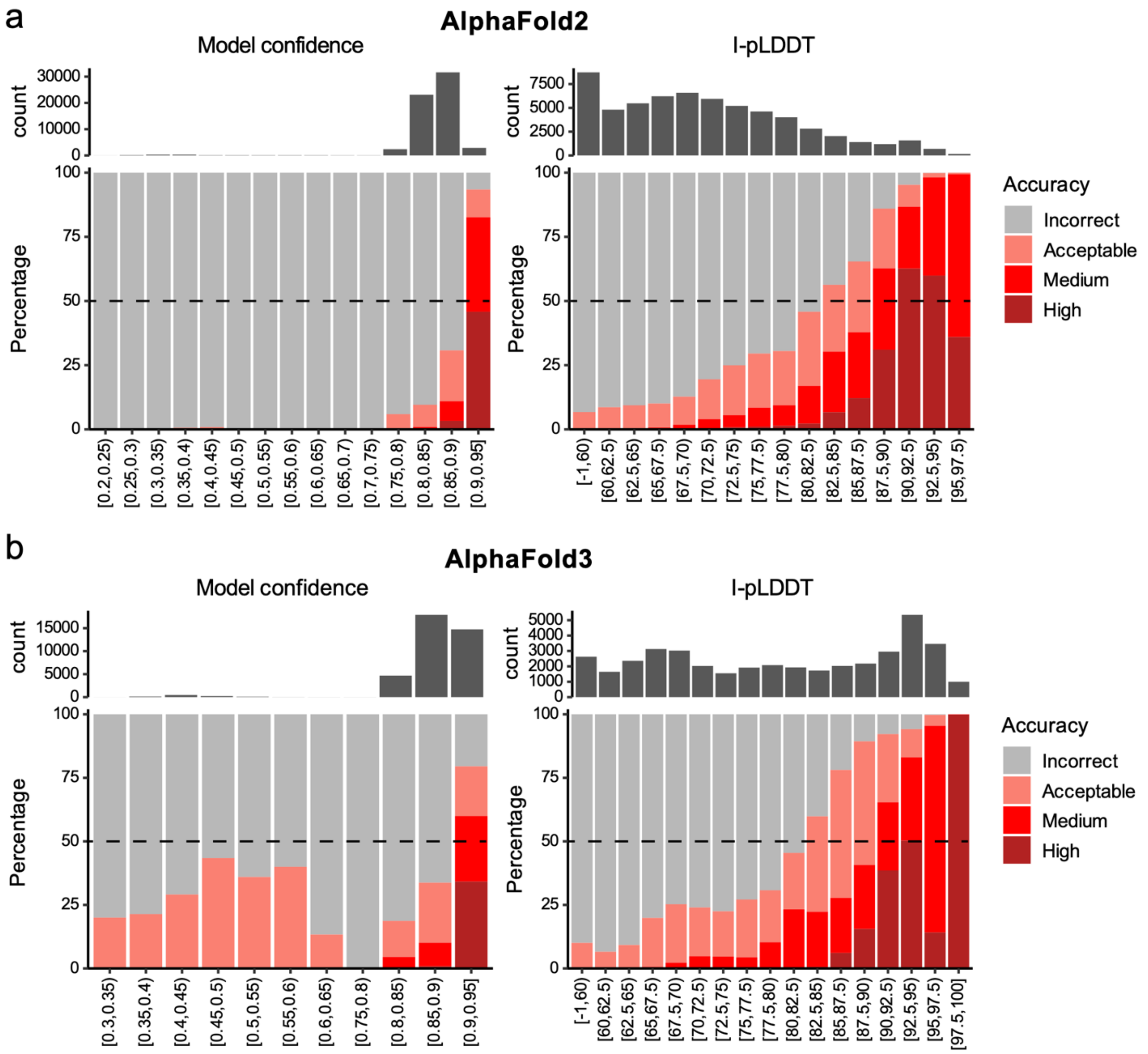


**Figure S5. Antibody-peptide success rates for model confidence and I-pLDDT score ranges.** (A) AlphaFold2 and (B) AlphaFold3 success rates for pooled antibody-peptide benchmark models are shown for model confidence (left) and I-pLDDT (right) score ranges, colored by CAPRI accuracy level as shown on right. Numbers of models per bin are shown at top, and dashed line at 50% is shown for reference.

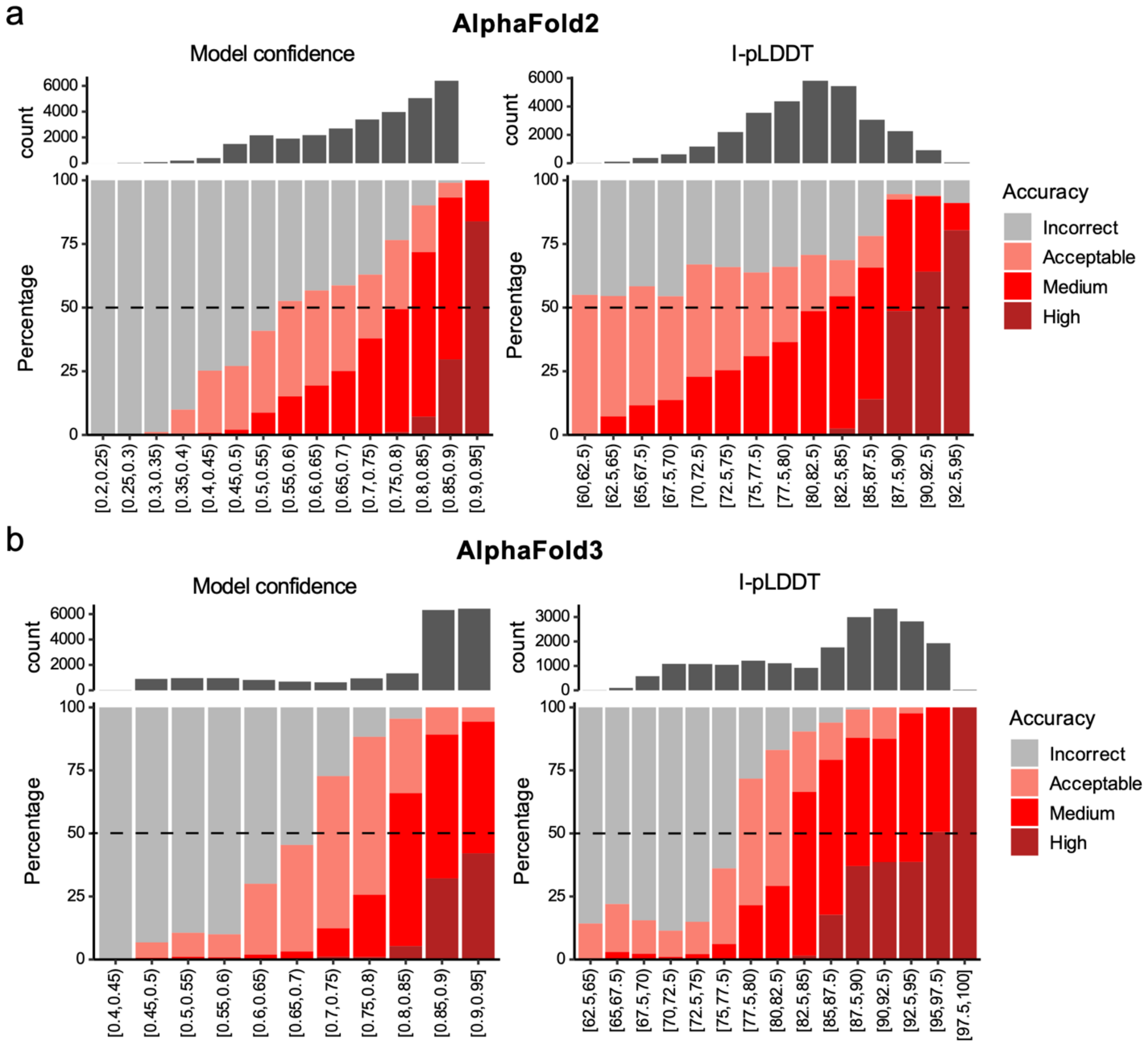


**Figure S6. TCR-pMHC success rates for model confidence and I-pLDDT score ranges.** (A) AlphaFold2 and (B) AlphaFold3 success rates for pooled TCR-pMHC benchmark models are shown for model confidence (left) and I-pLDDT (right) score ranges, colored by CAPRI accuracy level as shown on right. Numbers of models per bin are shown at top, and dashed line at 50% is shown for reference.

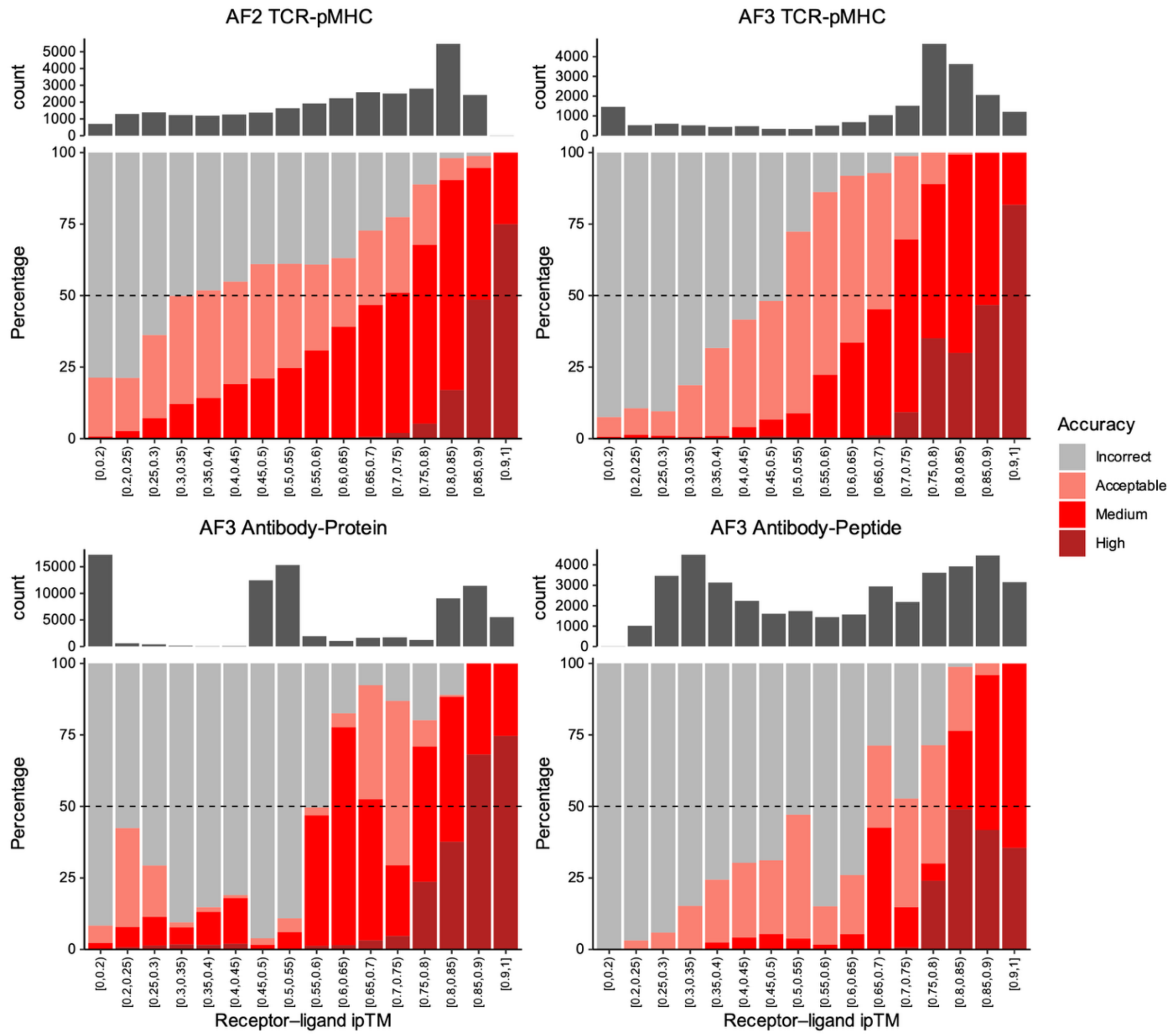


**Figure S7. Success rates for RL-ipTM score ranges.** Binned success percentage rates are shown for receptor-ligand ipTM (RL-ipTM) score ranges for TCR-pMHC benchmark models from TCRmodel2 (AF2; top left) and AF3 (top right), as well as antibody-protein and antibody-peptide AF3 models (bottom left and bottom right), colored by CAPRI accuracy levels as shown on right. Numbers of models per bin are shown at top, and dashed line at 50% is shown for reference.

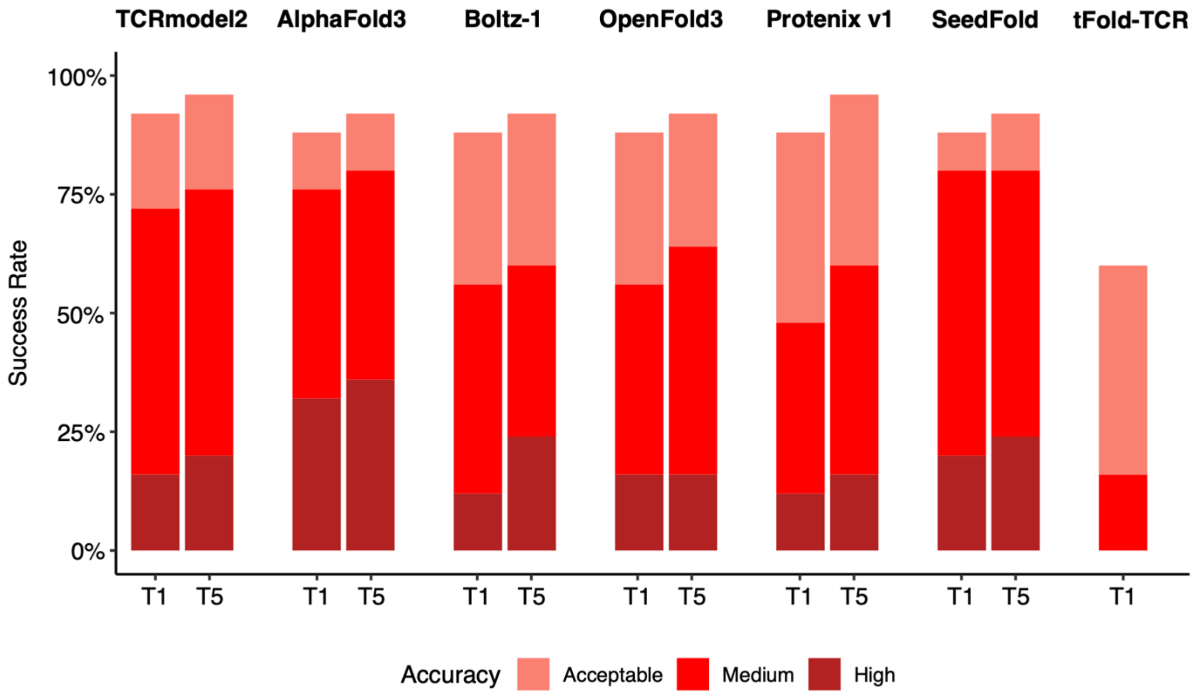


**Figure S8. TCR-pMHC success comparison for additional deep learning modeling methods.** Success rates for modeling TCR-pMHC benchmark cases (N=25) using TCRmodel2, AlphaFold3, Boltz-1, OpenFold3, Protenix v1, SeedFold, and tFold-TCR. Each method was run using one seed, generating five ranked models per complex, except for tFold-TCR which generates one model per complex. Predictions from each method were assessed for top-ranked (T1) and all model (T5, when available) success rates for CAPRI accuracy levels, represented by separate colors as shown on bottom.

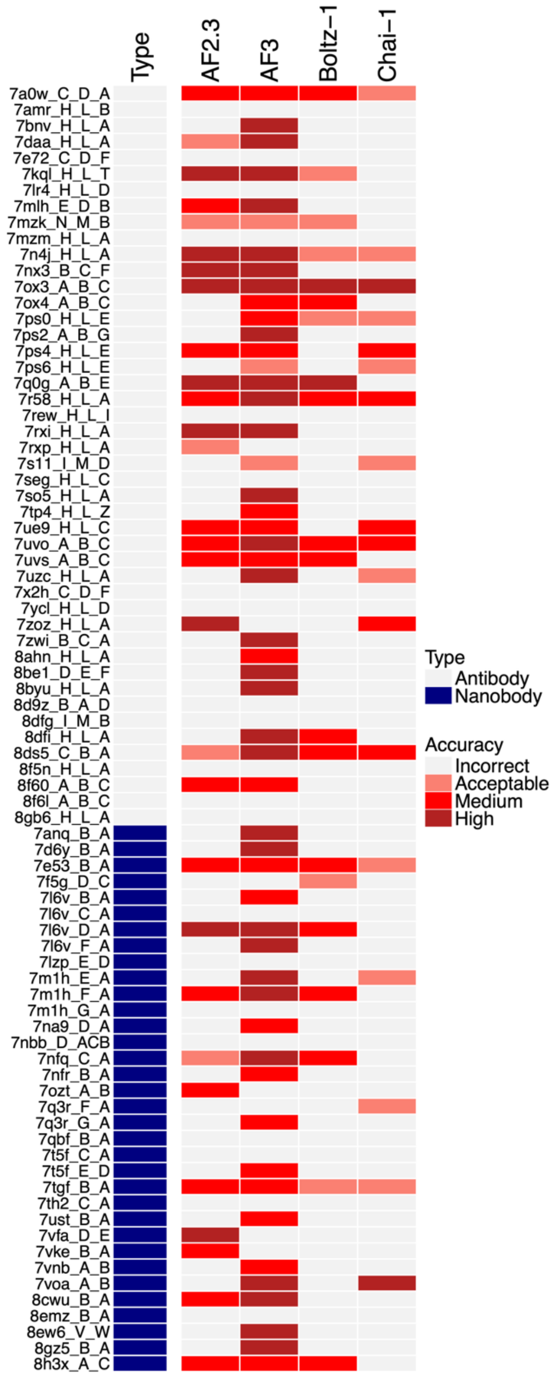


**Figure S9. Top-ranked model per-case success for antibody-protein complexes.** Top-ranked (T1) model accuracy for individual antibody-protein test cases for AF2.3, AF3, Boltz-1, and Chai-1 is shown, with accuracy levels colored as shown on right. Test cases are given by complex PDB code and chain IDs (separated by underscores; left) and antibody type (antibody, nanobody) is indicated by left column, colored as indicated on right. All methods generated 100 structures per complex, which were ranked by model confidence score to obtain the top-ranked model.

**Table S1.** Antibody-protein benchmark test cases.

| PDB[a] | Heavy[b] | Light[b] | Antigen[b] | Release Date[a] | Resolution, Å[a] |
|---|---|---|---|---|---|
| 7tgf | B | - | A | 2022-10-12 | 1.35 |
| 8emz | B | - | A | 2023-03-15 | 1.40 |
| 7kql | H | L | T | 2022-02-09 | 1.49 |
| 7nbb | D | - | A C B | 2022-08-10 | 1.55 |
| 8f60 | A | B | C | 2023-05-24 | 1.64 |
| 8h3x | A | - | C | 2023-02-08 | 1.66 |
| 7d6y | B | - | A | 2021-10-06 | 1.67 |
| 7nfq | C | - | A | 2021-12-01 | 1.68 |
| 7ox3 | A | B | C | 2022-12-28 | 1.70 |
| 7ust | B | - | A | 2022-12-28 | 1.70 |
| 8gz5 | B | - | A | 2022-12-07 | 1.70 |
| 8cwu | B | - | A | 2022-07-06 | 1.71 |
| 7ozt | A | - | B | 2022-07-13 | 1.74 |
| 7ue9 | H | L | C | 2022-06-08 | 1.75 |
| 8gb6 | H | L | A | 2023-05-24 | 1.75 |
| 7f5g | D | - | C | 2022-05-25 | 1.75 |
| 7vfa | D | - | E | 2022-03-30 | 1.75 |
| 7rxp | H | L | A | 2022-03-16 | 1.76 |
| 7na9 | D | - | A | 2021-12-22 | 1.76 |
| 7so5 | H | L | A | 2022-05-11 | 1.80 |
| 7ox4 | A | B | C | 2022-12-28 | 1.80 |
| 8d9z | B | A | D | 2023-06-14 | 1.80 |
| 8ahn | H | L | A | 2023-05-24 | 1.80 |
| 7voa | A | - | B | 2022-08-10 | 1.80 |
| 7q0g | A | B | E | 2021-12-22 | 1.82 |
| 7th2 | C | - | A | 2022-10-12 | 1.84 |
| 8byu | H | L | A | 2023-07-12 | 1.85 |
| 8f6l | A | B | C | 2023-05-24 | 1.85 |
| 7qbf | B | - | A | 2022-03-16 | 1.85 |
| 7nfr | B | - | A | 2021-12-01 | 1.88 |
| 8dfi | H | L | A | 2022-10-12 | 1.90 |
| 8f5n | H | L | A | 2023-03-08 | 1.90 |
| 7r58 | H | L | A | 2022-11-16 | 1.90 |
| 7zwi | B | C | A | 2022-06-22 | 1.90 |
| 8ew6 | V | - | W | 2022-11-23 | 1.90 |
| 7vke | B | - | A | 2022-08-03 | 1.90 |
| 7q3r | G | - | A | 2022-11-16 | 1.92 |
| 7q3r | F | - | A | 2022-11-16 | 1.92 |
| 8ds5 | C | B | A | 2022-09-28 | 1.93 |
| 7ps4 | H | L | E | 2021-12-15 | 1.94 |
| 7amr | H | L | B | 2022-04-20 | 1.95 |
| 7tp4 | H | L | Z | 2022-02-23 | 1.95 |
| 8be1 | D | E | F | 2023-01-25 | 1.98 |
| 8dfg | I | M | B | 2022-10-12 | 2.00 |

| | | | | | |
|---|---|---|---|---|---|
| 7l6v | B | - | A | 2021-12-22 | 2.01 |
| 7l6v | D | - | A | 2021-12-22 | 2.01 |
| 7l6v | C | - | A | 2021-12-22 | 2.01 |
| 7l6v | F | - | A | 2021-12-22 | 2.01 |
| 7a0w | C | D | A | 2022-02-23 | 2.04 |
| 7uvs | A | B | C | 2023-02-15 | 2.06 |
| 7e72 | C | D | F | 2021-11-10 | 2.09 |
| 7uvo | A | B | C | 2023-02-15 | 2.09 |
| 7x2h | C | D | F | 2023-03-01 | 2.10 |
| 7zoz | H | L | A | 2023-06-28 | 2.10 |
| 7mlh | E | D | B | 2022-05-04 | 2.10 |
| 7rew | H | L | I | 2022-05-25 | 2.10 |
| 7lr4 | H | L | D | 2021-12-15 | 2.10 |
| 7ycl | H | L | D | 2023-02-01 | 2.13 |
| 7rxi | H | L | A | 2022-03-16 | 2.15 |
| 7seg | H | L | C | 2021-11-24 | 2.16 |
| 7uzc | H | L | A | 2023-01-11 | 2.20 |
| 7anq | B | - | A | 2021-10-20 | 2.20 |
| 7e53 | B | - | A | 2021-10-13 | 2.21 |
| 7n4j | H | L | A | 2021-10-06 | 2.21 |
| 7mzk | N | M | B | 2021-10-06 | 2.25 |
| 7ps6 | H | L | E | 2021-12-15 | 2.26 |
| 7vnb | A | - | B | 2021-11-24 | 2.27 |
| 7mzm | H | L | A | 2021-10-06 | 2.30 |
| 7bnv | H | L | A | 2021-11-17 | 2.35 |
| 7daa | H | L | A | 2021-10-20 | 2.51 |
| 7s11 | I | M | D | 2021-11-03 | 2.58 |
| 7t5f | C | - | A | 2021-12-29 | 2.60 |
| 7t5f | E | - | D | 2021-12-29 | 2.60 |
| 7m1h | G | - | A | 2021-12-22 | 2.78 |
| 7m1h | E | - | A | 2021-12-22 | 2.78 |
| 7m1h | F | - | A | 2021-12-22 | 2.78 |
| 7nx3 | B | C | F | 2021-10-27 | 2.81 |
| 7lzp | E | - | D | 2021-12-22 | 2.86 |
| 7ps0 | H | L | E | 2021-12-15 | 2.92 |
| 7ps2 | A | B | G | 2021-12-15 | 2.99 |

[a]PDB code, resolution, and release date of complex structure
[b]Chain ID(s) of heavy, light, and antigen chains

**Table S2.** Antibody-peptide benchmark test cases.

| PDB[a] | Heavy[b] | Light[b] | Antigen[b] | Release date[a] | Resolution, Å[a] |
|---|---|---|---|---|---|
| 7y8j | H | L | A | 2023-07-12 | 1.03 |
| 7rpu | A | B | P | 2022-07-13 | 1.27 |
| 7doh | H | L | I | 2021-10-27 | 1.45 |
| 8d36 | H | L | F | 2022-07-27 | 1.45 |
| 8dtr | E | F | J | 2022-11-23 | 1.5 |
| 8op0 | A | - | B | 2023-05-31 | 1.54 |
| 8dtx | A | B | G | 2022-11-23 | 1.6 |
| 8bbh | H | L | A | 2022-12-21 | 1.62 |
| 7q4q | B | A | E | 2022-06-15 | 1.65 |
| 8duz | C | D | E | 2023-07-05 | 1.65 |
| 7u0a | H | L | A | 2022-08-03 | 1.7 |
| 7raq | H | L | P | 2022-04-13 | 1.74 |
| 8dtt | E | F | J | 2022-11-23 | 1.75 |
| 8f0l | A | B | Q | 2023-03-29 | 1.81 |
| 7skz | H | L | A | 2022-07-20 | 1.86 |
| 8dgu | H | L | A | 2023-01-25 | 1.89 |
| 7s3n | H | L | A | 2021-10-27 | 1.9 |
| 7rlx | A | B | P | 2022-01-26 | 1.97 |
| 7n08 | H | L | F | 2022-03-30 | 2 |
| 8d47 | H | L | C | 2022-12-21 | 2 |
| 7n0x | H | L | G | 2022-04-06 | 2 |
| 7tcq | H | L | C | 2022-08-03 | 2.02 |
| 7u0e | H | L | C | 2022-08-03 | 2.1 |
| 7u09 | H | L | A | 2022-08-03 | 2.1 |
| 8fax | A | B | L | 2023-05-03 | 2.1 |
| 7sjp | H | L | E | 2022-09-07 | 2.1 |
| 7s4g | E | F | I | 2022-04-13 | 2.2 |
| 7uym | H | L | P | 2022-11-23 | 2.2 |
| 7rqr | A | B | C | 2022-03-02 | 2.23 |
| 8d6z | A | B | H | 2022-07-27 | 2.3 |
| 8dgv | H | L | A | 2023-01-25 | 2.3 |
| 8fg0 | A | B | Q | 2023-05-10 | 2.36 |
| 7um3 | I | M | B | 2022-09-07 | 2.4 |
| 7k7r | B | A | C | 2022-01-12 | 2.5 |
| 7sl5 | A | B | C | 2022-07-20 | 2.5 |
| 7e6p | H | L | A | 2022-01-05 | 2.5 |
| 7x9e | C | D | F | 2022-05-11 | 2.6 |
| 7y3j | H | L | A | 2022-08-17 | 2.6 |
| 8dao | C | D | I | 2022-07-27 | 2.8 |
| 8dgw | C | D | J | 2023-01-25 | 2.81 |
| 8dgx | H | L | C | 2023-01-25 | 2.89 |

[a]PDB code, resolution, and release date of complex structure
[b]Chain ID(s) of heavy, light, and antigen chains

**Table S3.** TCR-pMHC benchmark test cases.

| PDB[a] | Release date[a] | Resolution[a] | TCR name | Class[b] |
|---|---|---|---|---|
| 6zkw | 2022-01-26 | 2.26 | InhA:01 | 1 |
| 7na5 | 2022-06-22 | 2.5 | 47BE7 | 1 |
| 7ow5 | 2022-07-20 | 2.58 | JDIa41b1 | 1 |
| 7qpj | 2022-08-03 | 1.54 | c756 | 1 |
| 8gvb | 2022-10-19 | 3.2 | TD08 | 1 |
| 8d5q | 2022-09-14 | 2.50 | TG6 | 1 |
| 7phr | 2022-08-31 | 3.08 | GPa3b17 | 1 |
| 7ndq | 2022-01-26 | 2.55 | Gag:02 | 1 |
| 7rrg | 2022-03-23 | 2.12 | 0606T1-2 | 1 |
| 7l1d | 2022-03-23 | 3.11 | 21LT2-2 | 1 |
| 7rk7 | 2022-11-02 | 2.54 | TIL 1383i | 1 |
| 7dzm | 2022-01-26 | 2.25 | T18A | 1 |
| 7n2n | 2022-12-07 | 2.6 | AS4.2 | 1 |
| 8gom | 2023-02-22 | 2.78 | RLQ7 | 1 |
| 7sg0 | 2022-02-23 | 3 | W316 | 2 |
| 7sg1 | 2022-02-23 | 3.1 | XPA5 | 2 |
| 7rdv | 2022-07-27 | 2.90 | TFH | 2 |
| 7z50 | 2022-07-20 | 2.65 | 4.1 | 2 |
| 7t2c | 2022-12-28 | 3.1 | B5 | 2 |
| 7t2b | 2022-12-28 | 2.8 | 5F | 2 |
| *8dnt* | *2023-07-19* | *3.18* | *LLL8* | *1* |
| *8en8* | *2024-03-27* | *2.70* | *3180* | *1* |
| *8f5a* | *2024-05-22* | *1.95* | *KS1* | *1* |
| *8qfy* | *2024-05-15* | *2.33* | *a42b20* | *1* |
| *8wte* | *2024-05-01* | *2.17* | *4TCR2* | *1* |

[a]PDB code, resolution, and release date of complex structure
[b]MHC class for the TCR-pMHC complex structure

**Table S4**. Proportions of CAPRI high and medium or higher accuracy models in pooled sets of AF2 and AF3 models used for confidence score analysis.

| | **AlphaFold2** | | **AlphaFold3** | |
|---|---|---|---|---|
| | **% High** | **% High/Med** | **% High** | **% High/Med** |
| Antibody-Protein | 2.8 | 13.4 | 19.8 | 38.2 |
| Antibody-Peptide | 3.8 | 9.8 | 14.1 | 33.0 |
| TCR-pMHC | 7.8 | 48.2 | 24.1 | 64.8 |

**Table S5**. AlphaFold2 and AlphaFold3 confidence score model accuracy classification performance (ROC AUC) and correlations with model accuracy (DockQ).

| | | High vs. Rest | | High/Medium vs. Rest | | DockQ PCC[a] | | DockQ SCC[a] | |
|---|---|---|---|---|---|---|---|---|---|
| **Complex Type** | **Score Type** | **AF2** | **AF3** | **AF2** | **AF3** | **AF2** | **AF3** | **AF2** | **AF3** |
| Antibody-Protein | Model confidence | 0.97 | 0.94 | 0.90 | 0.97 | 0.60 | 0.85 | 0.52 | 0.81 |
| | ipTM | 0.97 | 0.95 | 0.89 | 0.97 | 0.57 | 0.83 | 0.51 | 0.82 |
| | Ab-Ag ipTM | - | 0.94 | - | 0.97 | - | 0.79 | - | 0.80 |
| | I-pLDDT | 0.97 | 0.96 | 0.83 | 0.97 | 0.41 | 0.85 | 0.27 | 0.72 |
| Antibody-Peptide | Model confidence | 0.95 | 0.84 | 0.91 | 0.90 | 0.33 | 0.41 | 0.53 | 0.71 |
| | ipTM | 0.95 | 0.81 | 0.92 | 0.87 | 0.32 | 0.36 | 0.53 | 0.66 |
| | Ab-Ag ipTM | - | 0.88 | - | 0.95 | - | 0.81 | - | 0.81 |
| | I-pLDDT | 0.97 | 0.88 | 0.92 | 0.94 | 0.57 | 0.82 | 0.44 | 0.80 |
| TCR-pMHC | Model confidence | 0.91 | 0.78 | 0.87 | 0.92 | 0.65 | 0.85 | 0.68 | 0.73 |
| | ipTM | 0.91 | 0.79 | 0.87 | 0.93 | 0.65 | 0.85 | 0.68 | 0.76 |
| | TCR-pMHC ipTM | 0.92 | 0.82 | 0.86 | 0.96 | 0.64 | 0.85 | 0.68 | 0.78 |
| | I-pLDDT | 0.95 | 0.79 | 0.73 | 0.93 | 0.39 | 0.83 | 0.43 | 0.74 |

[a]Pearson correlation coefficient (PCC) or Spearman correlation coefficient (SCC) with model DockQ accuracy values for pooled set.

**Table S6**. AlphaFold2 and AlphaFold3 score cutoff values for 50% high accuracy models.

| | | AF2 | | AF3 | |
|---|---|---|---|---|---|
| **Complex Type** | **Score Type** | **Cutoff[a]** | **Recall[a]** | **Cutoff[a]** | **Recall[a]** |
| Antibody-Protein | Model confidence | 0.91 | 0.11 | 0.66 | 0.99 |
| | ipTM | 0.91 | 0.12 | 0.64 | 0.99 |
| | Ab-Ag ipTM | - | - | 0.62 | 1.00 |
| | I-pLDDT | 93 | 0.36 | 79 | 1.00 |
| Antibody-Peptide | Model confidence | 0.90 | 0.47 | 0.99 | 0.00 |
| | ipTM | 0.90 | 0.54 | 0.96 | 0.00 |
| | Ab-Ag ipTM | - | - | 0.91 | 0.18 |
| | I-pLDDT | 87 | 0.78 | 96 | 0.20 |
| TCR-pMHC | Model confidence | 0.89 | 0.07 | 0.94 | 0.44 |
| | ipTM | 0.88 | 0.06 | 0.92 | 0.37 |
| | TCR-pMHC ipTM | 0.85 | 0.47 | 0.84 | 0.43 |
| | I-pLDDT | 87 | 0.80 | 95 | 0.21 |

[a]Cutoff score for 50% positive predictive value (PPV; precision), and recall value at that score cutoff.

**Table S7**. Modeling accuracy for noncanonical TCR-pMHC complex (9EJG).

| **Protocol** | **Success** | **Model confidence** | **I-pLDDT** | **I-RMSD (Å)[a]** | **L-RMSD (Å)[a]** | **DockQ[a]** | **CAPRI[a]** |
|---|---|---|---|---|---|---|---|
| **AF3** | T1 | 0.892 | 89.9 | 14.42 | 29.1 | 0.030 | Incorrect |
| | T3 | 0.876 | 93.1 | 0.97 | 5.01 | 0.786 | High |
| | T100 | 0.874 | 93.0 | 0.84 | 3.68 | 0.852 | High |
| **TCRmodel2** | T1 | 0.865 | 92.1 | 20.98 | 64.91 | 0.007 | Incorrect |
| | T100 | 0.403 | 75.4 | 10.11 | 22.36 | 0.049 | Incorrect |
| **Boltz-1** | T1 | 0.721 | 79.8 | 24.03 | 68.17 | 0.006 | Incorrect |
| | T100 | 0.708 | 81.0 | 23.56 | 67.20 | 0.007 | Incorrect |
| **OpenFold3** | T1 | 0.875 | 81.2 | 24.28 | 69.94 | 0.006 | Incorrect |
| | T100 | 0.849 | 70.7 | 14.76 | 31.56 | 0.026 | Incorrect |
| **SeedFold** | T1 | 0.797 | 89.6 | 14.32 | 28.43 | 0.031 | Incorrect |
| | T100 | 0.494 | 80.9 | 14.30 | 26.13 | 0.036 | Incorrect |

Reported metrics represent the scores and accuracy of the top-ranked model (T1 success), the most accurate model out of the top 3 ranked models (T3 success; AF3 only), or the most accurate model out of all 100 models generated (T100 success), as noted in "Success" column.

[a]I-RMSD, L-RMSD, DockQ, and CAPRI accuracy levels were calculated by the DockQ program by comparison of models with native complex structure (PDB code 9EJG).
